# Stretching the structural envelope of isomeric imatinib analogs that reduce β-amyloid production by modulating both β- and γ-secretase cleavages of APP

**DOI:** 10.1101/2024.07.14.602669

**Authors:** William J. Netzer, Anjana Sinha, Mondana Ghias, Emily Chang, Katherina Gindinova, Emily Mui, Ji-Seon Seo, Subhash C. Sinha

## Abstract

We previously showed that the anticancer drug imatinib mesylate (IMT, trade name: Gleevec) and a chemically distinct compound, DV2-103 (a kinase-inactive derivative of the potent Abl and Src kinase inhibitor, PD173955) lower Aβ levels at low micromolar concentrations primarily through a lysosome-dependent mechanism that renders APP less susceptible to proteolysis by BACE1 without directly inhibiting BACE1 enzymatic activity, or broadly inhibiting the processing of other BACE1 substrates. Additionally, IMT indirectly inhibits γ-secretase and stimulates autophagy, and thus may decrease Aβ levels through multiple pathways. In two recent studies we demonstrated similar effects on APP metabolism caused by derivatives of IMT and DV2-103. In the present study we investigated how so many structurally diverse compounds affect APP metabolism in the same way, with similar potencies and production of APP metabolites. To this end, we synthesized and tested radically altered IMT regioisomers that possess medium structural similarity to IMT. Independent of structural similarity, these isomers manifest widely differing potencies in altering APP metabolism. These will enable us to choose the most potent isomers for further derivatization.

## 1 Introduction

Neurotoxic β-amyloid peptides (Aβ) are major drivers of Alzheimer’s disease (AD) and are formed by sequential cleavage of the amyloid precursor protein (APP) by β-secretase (BACE1/2) and γ-secretase, respectively. Both β- and γ-secretases can be pharmacologically inhibited to reduce production of Aβ peptides. Indeed, there has been great interest in the development of inhibitors and modulators of the secretases as potential AD therapeutics(Miranda et al. 2021; Kumar et al. 2018; Zhao et al. 2020; Portelius et al. 2010; Hur 2022; Panza et al. 2009; Golde et al. 2013; Rynearson et al. 2021) but at this time all clinical trials involving secretase inhibitors/modulators have failed. Reasons given have included timing of drug administration (too late in disease course for benefits to occur); non-specific inhibition of secretase substrates other than APP; lack of target engagement; toxicity; and even failure of the Amyloid hypothesis(Kim et al. 2022).

In our previous study, we have shown that the anticancer drug IMT, which is a potent Abl kinase inhibitor(Druker et al. 1996), and PD173955(Nagar et al. 2002), an Abl/Src kinase inhibitor, reduce Aβ production in cultured N2a695 cells, rat embryonic neurons, and in guinea pig brain in vivo by indirectly inhibiting γ-secretase processing of APP, while sparing γ-secretase processing of Notch1 in cellular assays(Netzer et al. 2003). In a recent study we further showed that a kinase inactive derivative of PD173955, DV2-103, as well as IMT, reduced Aβ levels in cells mainly by indirectly inhibiting BACE cleavage of APP(Netzer et al. 2017), adding to our earlier study suggesting that the Aβ-lowering effect of IMT and DV2-103 are not only Abl kinase-independent but also broadly kinase-independent and affect both γ-secretase and BACE processing of APP. Additionally, IMT and DV2-103 decrease levels of APP-βCTF and sAPPβ, and raise levels of APP- αCTF, as well as a 141 amino acid APP-CTF (C141), and a 9 kDa APP-CTF (all consistent with reduced BACE processing of APP) in N2a695 cells(Netzer et al. 2017). Remarkably, we showed that this pattern of APP metabolites induced by IMT and DV2-103, and some of their analogs is observed when N2a695 cells are treated with a general, active-site-directed BACE inhibitor(Netzer et al. 2017; Sun et al. 2019; Sinha et al. 2019). We also demonstrated that IMT does not inhibit BACE1 enzymatic activity in two *in vitro* BACE1 assays at concentrations up to 100μM or inhibit processing of several non-APP BACE substrates in cells(Netzer et al. 2017). Additionally, we demonstrated that these inhibitory activities of IMT and DV2-103 require acidified lysosomes and we provided a model suggesting that the effects of these drugs on APP metabolism were a result of their effects on lysosomes, which caused APP to undergo increased trafficking to lysosomes and spend less time in the amyloidogenic pathway where Aβ and its direct precursor, the APP-βCTF, are formed(Netzer et al. 2017).

To understand how so many structurally different compounds reduce levels of secreted Aβ in cells and have a characteristic effect on APP metabolite levels, we designed and synthesized a derivative of IMT, referred to here as IMT isomer **1** (Fig. 1A), as well as additional IMT isomers **2**- **3** (Fig. 1E) and tested their effects on APP metabolism by measuring the Aβ40 levels in cell supernatants. We found that isomer **2**, not **3**, showed effects comparable to IMT and isomer **1** (See Fig. 1 for the activity data of IMT and isomer **1** and Table 1 isomers **2**-**3**). Subsequently, we focused on derivatives of isomer **1** and **2** and tested their effects on APP metabolism. The results are shown in Table 1. Overall, our goal was to make a large change in the structure of IMT that would greatly alter the pharmacophore structurally but still maintain IMT’s physical properties, in particular its property as a weak base, which is necessary for its sequestration in lysosomes through ion trapping(Kazmi et al. 2013; Burger et al. 2015).

**Fig. 1.**
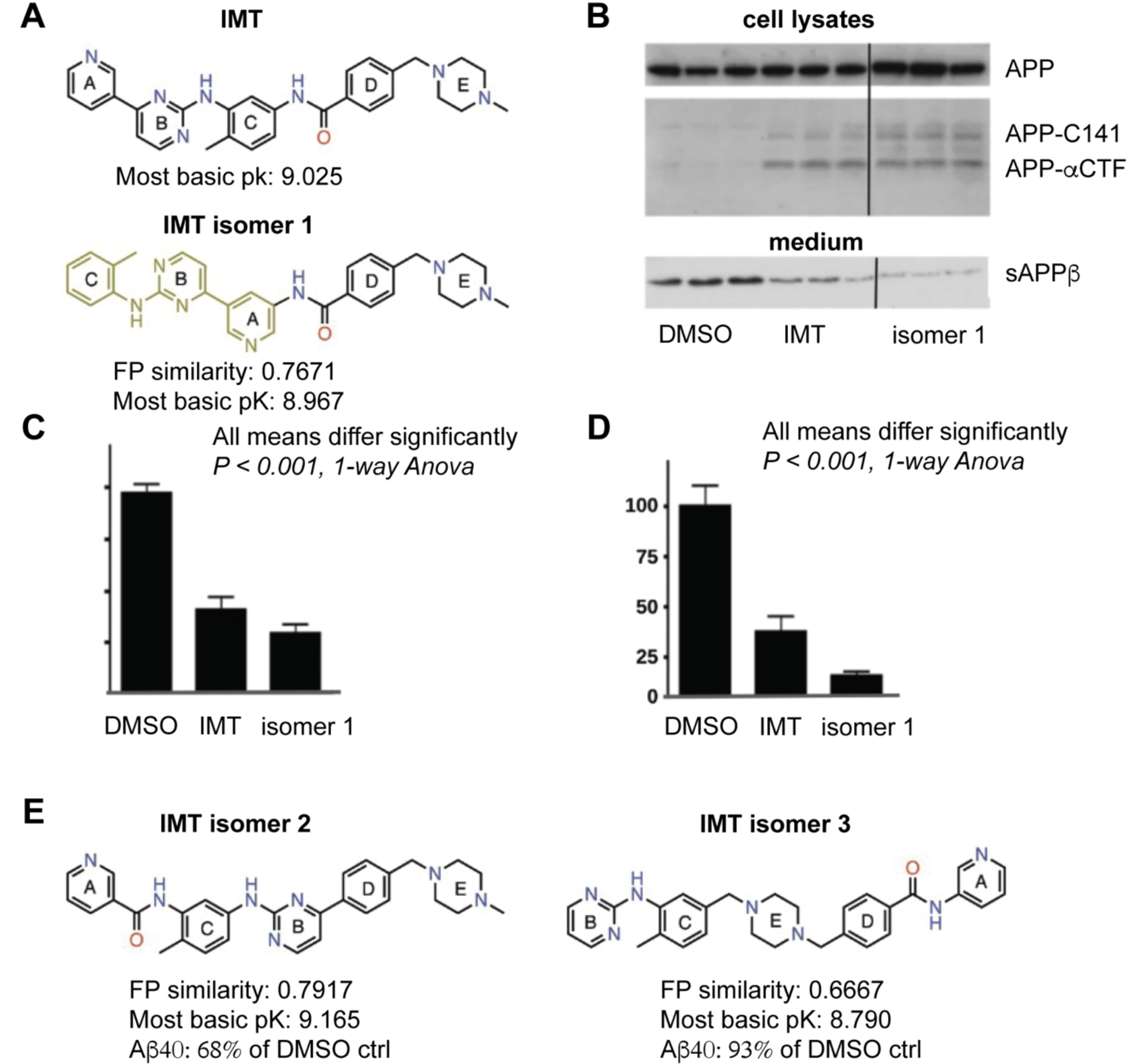
A Major Structural Change in IMT, results in IMT isomer 1, which retains IMT’s Aβ-reducing effect and its reduction of BACE processing of APP in N2a 695 cells. (A, E) Structures and properties of IMT and isomers **1**, **2** and **3**. Fingerprint (FP) similarity of *iso*-IMT **1**-**3** to parent IMT and their properties, including most basic PK, were calculated *in silico.* B) Western blots of N2a cell lysates (upper) and cell media (lower) from experiments using IMT and isomer **1** probed with antibody RU369 (anti-C terminal APP) and RU anti-C-terminal sAPPβ (bottom), respectively. Each western blot panel shows lanes from a single gel. However, the three lanes at the right of each, which refer to incubation with isomer **1**, are from a different part of the same gel. C) Quantification of secreted Aβ40 in N2a cells incubated with IMT or isomer **1**, 1-way Anova, p < 0.001. D) Quantification of sAPPβ levels. 1-way Anova, p < 0.001.

**Table 1.**
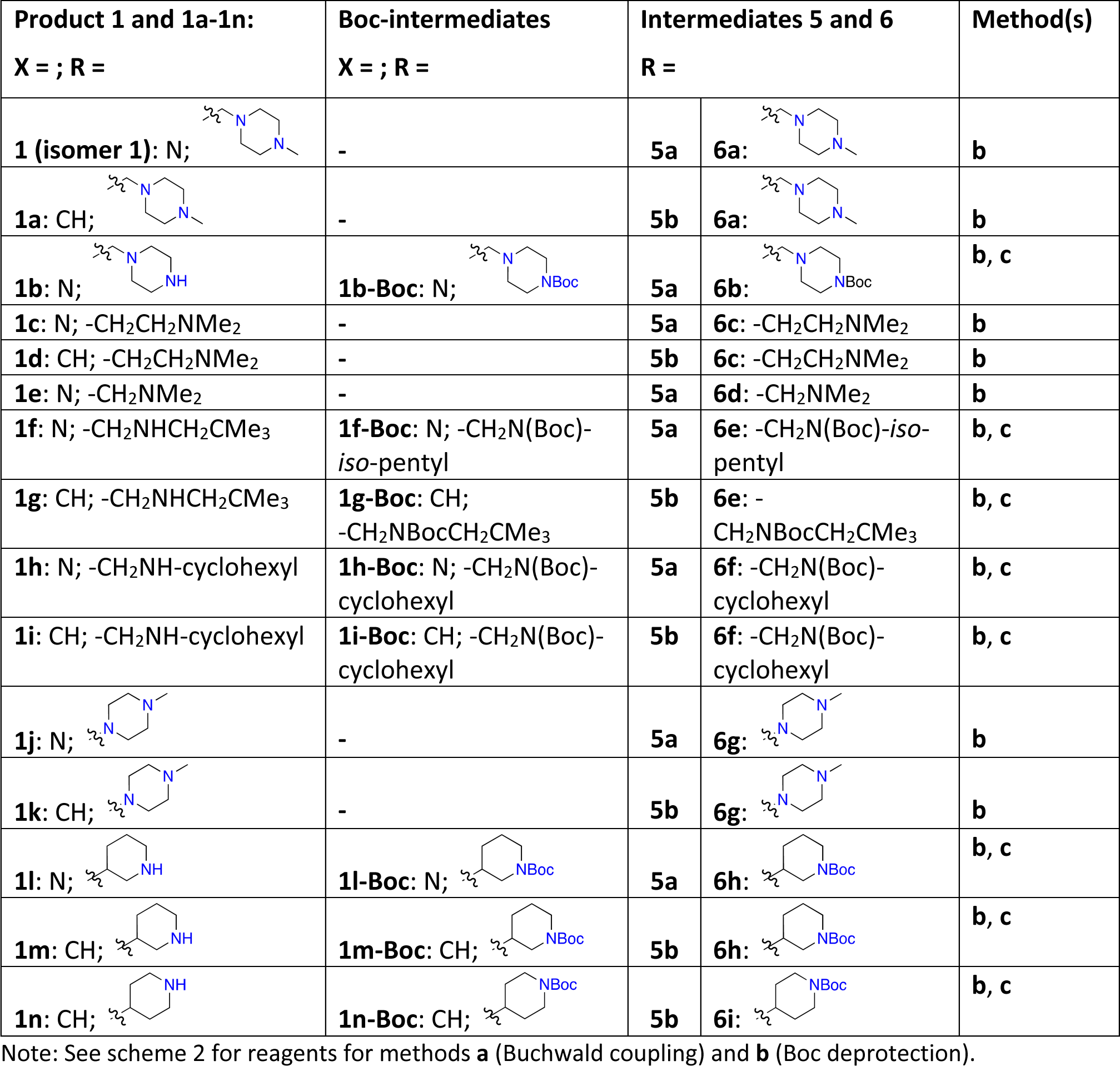
Intermediates (5a-5b, 6a-6i), step(s), and methods used to prepare IMT isomer 1 and 1a-1n.

## 2 Material and Methods

All commercial chemicals and solvents were reagent grade and used without further purification. All air-sensitive reactions were performed under argon protection. Column chromatography was performed using 230-400 mesh silica gel. Analytical thin layer chromatography was performed on 250 μM silica gel F_254_ plates. Preparative thin layer chromatography was performed on 1000 μM silica gel F_254_ plates. All final compounds were purified using HPLC. The identity and purity of each product was determined using MS, HPLC, TLC, and NMR analyses. ^1^H NMR spectra were recorded on either a Bruker 400 or 600 MHz instrument. Chemical shifts are reported in δ values in ppm downfield from TMS as the internal standard. ^1^H data are reported as follows: chemical shift, multiplicity (s = singlet, d = doublet, t = triplet, q = quartet, br = broad, m = multiplet), coupling constant (Hz), integration. Purity of target compounds has been determined to be >95% by LC/MS on a Waters autopurification system with PDA, MicroMass ZQ and ELSD detector and a reverse phase column (Waters X-Bridge C18, 4.6 x 150 mm, 5 µm) eluted with water/acetonitrile gradients, containing 0.1% TFA. All compounds tested in this study were prepared in house and their structures were confirmed using ^1^H NMR and MS analyses (Spectral data provided for new compounds only). Yields were not optimized. All final compounds were obtained in >95% purity as judged by LCMS analysis.

N2a695 were cultured in 1:1 OptiMem Reduced Serum Media (Life Technologies): Dulbecco’s Modified Eagle Medium ([+] 4.5 g/L D-glucose; [+] L-Glutamine; [-] Sodium pyruvate (Life Technologies) supplemented with 5% fetal bovine serum, 0.4% Penstrep and 0.4% Geneticin and incubated at 37 °C in 5% CO_2_. Antibodies were obtained from The Laboratory of Molecular and Cellular Neuroscience at The Rockefeller University. Human Aβ40 and Aβ42 ELISA plates (Life Technologies) and Plus MSD (Mesoscale Discovery) plates for Aβ Peptide (Aβ38, Aβ40 and Aβ42) Panel 1 (6E10) Kit (Catalog number K15200G) were obtained from Thermo Fisher, Life Technologies and Meso Scale Discovery.

### 2.1. Synthesis of IMT isomer 1 and analogs 1a-t

Prepared using intermediates **5a**-**c** (Scheme 2).

#### I) Intermediates 5a-c

To prepare intermediate **5a**, a solution of **4a** (800 mg, 2.96 mmol) and o-toluidine (1 ml, 9.29 mmol) in *i*-PrOH (6 mL) was added 1N HCl (3 mL), and the mixture was heated at 125 °C using Microwave for 2 h. Solvents were removed under reduced pressure, residues treated with aqueous NaHCO_3_ to neutralize, and the resulting mixture extracted with EtOAc. The combined organic layers were washed with brine, dried over anhydrous MgSO_4_, concentrated, and purified by Combi-Flash over Silica gel column using hexanes-EtOAc as eluants to afford intermediate **5a**. ^1^H NMR (600 MHz, CDCl_3_) of **5a**: δ 9.16 (s, 1H), 8.85 (s, 1H), 8.51 (s, 2H), 8.03 (d, *J* = 7.92 Hz, 1H), 7.35-7.30 (m, 2H), 7.14-7.07 (m, 2H), 2.38 (s, 3H); HRMS: *m/z* 341.0357 and 343.0337 [M+H]^+^, calcd. for C_16_H_14_BrN_4_; found 341.0391 and 343.0370.

Similarly, intermediate **4b** was reacted with o-toluidine using the method described for **5a** to afford intermediate **5b**, and **4c** reacted with 3-aminopyridine to afford **5c**. ^1^H NMR: (600 MHz, CDCl_3_) of **5b**: δ 8.47 (d, *J* = 6.0 Hz, 1H), 8.22 (s, 1H), 8.09 (d, *J* = 6.0 Hz, 1H), 7.96 (d, *J* = 6.0 Hz, 1H), 7.62 (d, *J* = 6.0 Hz), 7.36 (t, *J* = 6.0 Hz, 1H), 7.29 (t, *J* = 6.0 Hz, 2H), 7.24 (d, *J* = 12 Hz, 1H), 7.11 (d, *J* = 6.0 Hz, 1H), 7.06 (t, *J* = 6.0 Hz, 1H), 6.95 (s, 1H), 2.37 (s, 3H); HRMS: *m/z* 340.0439 and 342.0418, Calcd. for C_17_H_14_BrN_3_. MS of **5c**: *m/z* 264.12 [M+H]^+^.

#### II) Compounds 1 and 1a-n

To a degassed mixture of intermediates **5a** (38 mg) and **6a** (21 mg) in dioxane was added Pd_2_(dba)_3_ (4 mg), XanthPhos (7 mg), and Cs_2_CO_3_ (55 mg) and the resulting mixture was heated at 100°C temperature for 16 hours. Solvents were removed and worked up using EtOAc and water. Combined organic layers were dried over anhydrous MgSO_4_ and concentrated under reduced pressure. The resulting residues were purified by preparative TLC (Silica gel, 1 mm plate; CH_2_Cl_2_:MeOH:Aq. NH_3_ (90:10:1)) to afford the target product **1**. ^1^H NMR (400 MHz, CDCl_3_): d 9.01 (d, *J* = 1.5 Hz, 1H), 8.92 and 8.88 (s, 1H each), 8.47 (d, *J* = 5.12 Hz, 1H), 8.05 (d, *J* = 7.96 Hz, 1H), 7.88 (d, *J* = 7.96 Hz, 2H), 7.45 (d, *J* = 8.40 Hz, 1H), 7.28 (t, *J* = 4.28 Hz, 1H), 7.23 (d, *J* = 7.40 Hz, 1H), 7.15 (d, *J* = 5.16 Hz, 1H), 7.07-7.03 (m, 2H), 3.57 (s, 2H), 2.49 (br s, 8H), 2.35 and 2.30 (s, 3H each); HRMS: *m/z* 494.2624 [M+H]^+^, calcd. for C_29_H_32_N_7_O; found: 494.2653. Purity (HPLC): >98%.

#### Compound 1a

Buchwald coupling of **5b** with **6a** was performed as described for compound **1** to afford **1a**. ^1^H NMR (400 MHz, CDCl_3_) of **1a**: δ 8.46 (d, *J* = 5.20 Hz, 1H), 8.36 (s, 1H), 8.12 (d, 8.04 Hz, 1H), 8.02 (s, 1H), 7.87-7.82 (m, 4H), 7.52-7.46 (m, 3H), 7.31-7.23 (m, 1H), 7.17 (d, *J* = 5.16 Hz, 1H), 7.06 (t, *J* = 3.6 Hz, 1H), 6.96 (s, 1H), 3.56 (s, 2H), 2.52 (br s, 8H), 2.37 and 2.33 (s, 3H each); HRMS: *m/z* 493.2671 [M+H]^+^, calcd. for C_30_H_33_N_6_O; found: 493.2716.

#### Compound 1b

Intermediate **5a** reacted with **6b** under Buchwald conditions as described above for **1** to give the Boc protected compound **1b**-**Boc**. MS: *m/z* 579.30. To a solution of **1b-Boc** in EtOAc (3 mL) was added 4M HCl in dioxane (1 mL) at room temperature (RT) and the mixture was stirred for 2 hours or until the reaction was complete (monitored by TLC or LCMS). Solvents were removed under reduced pressure and purified over Silica gel column using CH_2_Cl_2_- MeOH/aq. NH_3_ to give the Boc-deprotected compound **1b**. HRMS: *m/z* 480.2467 [M+H]^+^, calcd. for C_28_H_30_N_7_O; found: 480.2506.

#### Compound 1c

Intermediate **5a** underwent Buchwald coupling with **6c** as described above for **1** to give compound **1c**. ^1^H NMR (600 MHz, CDCl_3_): δ 9.03, 8.95 and 8.90 (s, 1H each), 8.50 (d, *J* = 6.0 Hz, 1H), 8.09 (d, *J* = 6.0 Hz, 1H), 7.87 (d, *J* = 6.0 Hz, 2H), 7.35 (d, *J* = 6.0 Hz, 2H), 7.32-7.29 (m, 1H), 7.25 (d, *J* = 6.0 Hz, 1H), 7.20 (d, *J* = 6.0 Hz, 1H), 7.08 (d, *J* = 6.0 Hz, 2H), 2.89 (t, *J* = 6.0 Hz, 2H), 2.59 (d, *J* = 6.0 Hz, 2H), 2.37 (s, 3H), 2.33 (s, 6H); HRMS: *m/z* 453.2358 [M+H]^+^, calcd. for C_27_H_29_N_6_O; found: 453.2403.

#### Compound 1d

Intermediate **5b** underwent Buchwald coupling with **6c** as described for **1** to give compound **1d**. ^1^H NMR (600 MHz, CDCl_3_): δ 8.49 (d, *J* = 6.0 Hz, 1H), 8.38 (s, 1H), 8.15 (d, *J* = 6.0 Hz, 1H), 7.99 (br s, 1H), 7.88-7.83 (m, 4H), 7.68 (d, *J* = 6.0 Hz, 1H), 7.53 (t, *J* = 6.0 Hz, 1H), 7.42-7.38 (m, 2H), 7.33 (d, *J* = 12.0 Hz, 1H), 7.26 (d, *J* = 6.0 Hz, 1H), 7.21 (d, *J* = 6.0 Hz, 1H), 7.08 (t, *J* = 6.0 Hz, 1H), 6.97 (s, 1H), 2.29 (q, *J* = 6.0 Hz, 2H), 2.61 (q, *J* = 6.0 Hz, 2H), 2.40 (s, 3H), 2.38 (s, 6H); HRMS: *m/z* 452.2406 [M+H]^+^, calcd. for C_28_H_30_N_5_O; found 452.2450. Purity (HPLC): >98%.

#### Compound 1e

Intermediate **5a** underwent Buchwald coupling with **6d** as described for **1** to give compound **1e**. HRMS: *m/z* 439.2202 [M+H]^+^, calcd. for C_26_H_27_N_6_O; found: 439.2243.

#### Compound 1f

Intermediate **5a** underwent Buchwald coupling with **6e** as described for **1** to give the Boc-protected derivative, **1f-Boc**, and the latter underwent Boc-deprotection to give compound **1f**. ^1^H NMR (400 MHz, CDCl_3_) of **1f-Boc**: δ 9.04 (s, 1H), 8.43 (s, 1H), 8.85 (s, 1H), 8.49 (d, *J* = 5.04 Hz, 1H), 8.07 (d, *J* = 8.0 Hz, 1H), 7.89-7.87 (br, 2H), 7.27-7.17 (m, 6H), 7.07-7.03 (m, 2H), 4.58 (br, 2H), 3.16(br s, 1H), 3.07 (br s, 1H), 2.36 (s, 3H), 1.50 and 1.34 (s, 6H and 3H); 0.97 (s, 9H); MS: *m/z* 581.32 [M+H]^+^. ^1^H NMR (400 MHz, CDCl_3_) of **1f**: δ 9.06, 8.94 and 8.85 (s, 1H each), 8.50 (d, *J* = 5.04 Hz, 1H), 8.10 (d, *J* = 8.0 Hz, 1H), 8.0 (s, 1H), 8.07 (d, *J* = 8.0 Hz, 1H), 7.89 (d, *J* = 7.92 Hz, 2H), 7.52 (d, *J* = 7.80 Hz, 2H), 7.24-7.21 (m, 4H), 7.08 (t, J = 4.00 Hz, 1H), 7.00 (s, 1H), 3.92 (s, 2H), 2.38 (s, 2H), 2.37 (s, 3H), 0.95 (s, 9H); HRMS: *m/z* 481.2671 [M+H]^+^, calcd.; for: C_29_H_33_N_6_O; found 481.2716.

#### Compound 1g

Intermediate **5b** underwent Buchwald coupling with **6e** as described for **1** to give the Boc-protected derivative, **1g-Boc**, and the latter underwent Boc-deprotection to give compound **1g**. MS of **1g-Boc**: *m/z* 580.33 [M+H]^+^. ^1^H NMR (600 MHz, CDCl_3_) of **1g**: δ 8.47, 8.36 and 8.14 (s, 1H each), 7.88-7.83 (m, 4H), 7.50 (d, *J* = 7.40 Hz, 4H), 7.24-7.19 (m, 3H), 7.08--7.01 (m, 2H), 3.90 (s, 2H), 3.49 (s, 2H), 2.34 (s, 3H), 0.95 (m, 9H); HRMS: *m/z* 480.2719 [M+H]^+^, calcd. for C_30_H_34_N_5_O; found: 480.2776.

#### Compound 1h

Intermediate **5a** underwent Buchwald coupling with **6f** as described for **1** to give the Boc-protected derivative, **1h-Boc**, and the latter underwent Boc-deprotection to give compound **1h**. ^1^H NMR (400 MHz, CDCl_3_) of compound **1h-Boc**: δ 9.04 (br s, 1H), 8.97 and 8.88 (s, 1H each), 8.72 (br, 1H), 8.47 (d, *J* = 4.28 Hz, 1H), 8.06 (d, *J* = 7.92 Hz, 4H), 7.94 (m, 2H), 7.44 7.36-7.17 (m, 5H), 7.06 (m, 2H), 4.42 (br s, 2H), 4.08 (m, 1H), 2.36 (s, 3H), 1.70-1.4 (m, 8H), 1.35-1.26 (m, (9H+2H); MS: *m/z* 593.32 [M+H]^+^. ^1^H NMR (400 MHz, CDCl_3_) of compound **1h**: δ 9.03 (br s, 1H), 8.94 and 8.85 (s, each 1H), 8.48 (d, *J* = 4.84 Hz, 1H), 8.34 9 (br, 1H), 8.08 (d, *J* = 7.96 Hz, 1H), 7.88 (d, J = 7.80 Hz, 2H), 7.49 (d, *J* = 7.64 Hz, 2H), 7.30 (d, *J* = 7.56 Hz, 2H), 7.25-7.23 (m, 3H), 7.18 (d, *J* = 4.72 Hz, 1H), 7.09-7.05 (m, 2H), 3.94 (s, 2H), 2.57 (m, 1H), 2.37 (s, 3H), 1.98-1.45 (m, 8H), 1.29-1.18 (m, 2H); HRMS: *m/z* 493.2671 [M+H]^+^, calcd. for: C_30_H_33_N_6_O; found: 493.2718.

#### Compound 1i

Intermediate **5b** underwent Buchwald coupling with **6f** as described for **1** to give the Boc-protected derivative, **1i-Boc**, and the latter underwent Boc-deprotection to give compound **1i**. MS of **1i-Boc**: *m/z* 592.33 [M+H]^+^. ^1^H NMR (600 MHz, CDCl_3_) of **1i**: δ 8.41 (d, *J* = 5.04 Hz, 1H), 8.31 (s, 1H), 8.04 (d, *J* = 7.96 Hz, 1H), 7.88 (d, *J* = 7.72 Hz, 3H), 7.78 (d, *J* = 7.64 Hz, 1H), 7.49-7.42 (m, 4H), 7.27-7.24 (m, 3H), 7.15 (d, *J* = 4.96 Hz, 1H), 7.03 (t, *J* = 7.24 Hz, 1H), 3.90 (s, 2H), 3.38 (m, 1H), 2.34 (s, 3H), 1-97-1.62 (m, 6H), 1.26-1.16 (m, 4H); HRMS: *m/z* 492.2719 [M+H]^+^, calcd. for C_31_H_34_N_5_O; found 492.2772.

#### Compound 1j

Intermediate **5a** underwent Buchwald coupling with **6g** as described for **1** to give compound **1j**. ^1^H NMR (600 MHz, CDCl_3_): δ 9.04 (s, 1H), 8.94 (s, 1H), 8.86 (d, *J* = 6.0 Hz, 1H), 8.13 (dd, *J* = 12.0, 6.0 Hz, 1H), 7.87 (d, *J* = 6.0 Hz, 2H), 7.32 (t, *J* = 6.0 Hz, 2H), 7.265 (d, *J* = 6.0 Hz, 1H), 7.215 (d, *J* = 6.0 Hz, 1H), 7.08 (t, *J* = 6.0 Hz, 1H), 6.975 (d, *J* = 6.0 Hz, 2H), 3.40 (t, *J* = 6.0 Hz, 4H), 2.62 (t, *J* = 6.0 Hz, 4H), 2.40 and 2.39 (s, 3H each); MS: *m/z* 480.25 [M+H]^+^, calcd. for C_28_H_30_N_7_O; found: 480.25.

#### Compound 1k

Intermediate **5b** underwent Buchwald coupling with **6g** as described for **1** to give compound **1k**. ^1^H NMR (600 MHz, CDCl_3_) of **1k**: δ 8.49 (d, *J* = 6.0 Hz, 1H), 8.36 (s, 1H), 8.17 (dd, *J* = 6.0, 12.0 Hz, 1H), 7.88-7.83 (m, 3H), 7.51 (t, *J* = 6.0 Hz, 1H), 7.33-7.29 (m, 2H), 7.27 (d, *J* = 6.0 Hz), 7.22 (d, *J* = 6.0 Hz, 1H), 7.09 (t, *J* = 6.0 Hz, 1H), 6.99 (d, *J* = 6.0 Hz, 2H), 3.41 (t, *J* = 6.0 Hz, 4H), 2.63 (br t, 4H), 2.41 and 2.40 (s, 3H each); MS: *m/z* 479.25 [M+H]^+^, calcd. for C_28_H_31_N_6_O; found: 479.25.

#### Compound 1l

Intermediate **5a** underwent Buchwald coupling with **6h** as described for **1** to give the Boc-protected derivative, **1l-Boc**, and the latter underwent Boc-deprotection to give compound **1l**. ^1^HNMR (600 MHz, CDCl_3_) of **1l-Boc**: δ 9.04 (s, 1H), 8.95 (s, 1H), 8.88 (s, 1H), 8.50 (s, 2H), 8.10 (d, *J* = 6.0 Hz, 1H), 7.91 (d, *J* = 6.0 Hz, 2H), 7.37 (br t, J =, 2H), 7.30 (d, *J* = 6.0 Hz, 1H), 7.26 (d, *J* = 6.0 Hz, 1H), 7.20 (m, 2H), 4.18 (m, 1H), 2.82 (br, 2H), 2.40 (s, 3H), 2.08 (m, 1H), 1.82 (m, 1H), 1.69 (m, 2H), 1.61 (s, 3H), 1.49 (s, 6H), 1.48 (m, 2H); MS (ESI) *m/z* 564.28 [M]^+^, calcd. for C_33_H_36_N_6_O_3_; found: 565.29 [M+H]^+^. HRMS of **1l**: *m/z* 465.2358 [M+H]^+^, calcd. for C_28_H_29_N_6_O; found 465.2401.

#### Compound 1m

Intermediate **5b** underwent Buchwald coupling with **6h** as described for **1** to give the Boc-protected derivative, **1m-Boc**, and the latter underwent Boc-deprotection to give compound **1m**. ^1^HNMR (600 MHz, CDCl_3_) of **1m-Boc**: δ 8.50 (d, *J* = 6.0 Hz, 1H), 8.37 (s, 1H), 8.16 (d, *J* = 12.0 Hz, 1H), 7.93 (s, 1H), 7.89-7.86 (m, 3H), 7.545 (t, *J* = 6.0 Hz, 1H), 7.42 (d, *J* = 6.0 Hz, 1H), 7.32 (t, *J* = 12.0 Hz, 2H), 7.27 (d, *J* = 6.0 Hz, 1H), 7.22 (d, *J* = 6.0 Hz, 1H), 7.08 (t, *J* = 6.0 Hz, 1H), 6.96 (s, 1H), 4.18 (m, 1H), 2.81 (br, 2H), 2.40 (s, 3H), 2.08 (m, 1H), 1.81 (m, 1H), 1.68 (m, 2H), 1.59 (s, 3H), 1.51 (s, 6H), 1.48 (m, 2H); MS: *m/z* 564.29 [M+H]^+^. HRMS of **1m**: *m/z* 464.2406 [M+H]^+^, calcd. for C_29_H_29_N_5_O; found 464.2459. Purity (HPLC): >98%.

#### Compound 1n

Intermediate **5b** underwent Buchwald coupling with **6i** as described for **1** to give the Boc-protected derivative, **1n-Boc**, and the latter underwent Boc-deprotection to give compound **1n**. MS of **1n-Boc**: *m/z* 564.29 [M+H]^+^. MS of **1n**: *m/z* 463.24 [M+H]^+^, calcd. for C_29_H_30_N_5_O; found 464.24.

#### Compound 1o

NaCNBH_3_ (3 equiv.) and AcOH (1 equiv.) were added sequentially to a solution of amine **1l** (1 equiv.) and paraformaldehyde (3 equiv.) in MeOH (3-5 mL/mmol) at ice-water temperature and the reaction mixture was stirred at RT for another 2-8 hrs. Solvents were removed under reduced pressure, and the residue was suspended in CH_2_Cl_2_ and washed using water. Combined organic layers were dried using Na_2_SO_4_, filtered, and concentrated under reduced pressure. The residue was purified by Silica gel column to afford **1o**. ^1^H NMR (600 MHz, CD_3_OD+CDCl_3_): δ 8.99 (br s, 1H), 8.94 (s, 1H), 8.45 (s, 1H), 8.02 (d, *J* = 6.0 Hz, 2H), 7.74 (d, *J* = 6.0 Hz, 1H), 7.49 (d, *J* = 6.0 Hz, 2H), 7.30-7.24 (m, 3H), 7.10 (t, *J* = 6.0 Hz, 1H), 3.49 (m, 2H), 3.10 (t, *J* = 12.0 Hz, 1H), 2.95 (t, *J* = 12.0 Hz, 1H), 2.87 (m, 1H), 2.81 (s, 3H), 2.34 (s, 3H), 2.10-1.64 (m, 4H); HRMS: *m/z* 479.2515 [M+H]^+^, calcd. for C_29_H_31_N_6_O; found: 479.2560.

#### Compound 1p

Prepared by reductive amination of amine **1m** with paraformaldehyde and NaCNBH_3_ as described for **1o**. ^1^HNMR (600 MHz, CDCl_3_) of **1p**: δ 8.42 (d *J* = 6.0 Hz, 1H), 8.36 (s, 1H), 8.05 (d, *J* = 6.0 Hz, 1H), 7.87 (d, *J* = 6.0 Hz, 2H), 7.80 (d, *J* = 6.0 Hz, 1H), 7.47 (t, *J* = 6.0 Hz, 1H), 7.34 (d, *J* = 12.0 Hz, 2H), 7.265 (d, *J* = 6.0 Hz, 1H), 7.23 (d, *J* = 6.0 Hz, 1H), 7.16 (q, *J* = 6.0 Hz, 1H), 7.05 (t, *J* = 6.0 Hz, 1H), 3.29 (d, *J* = 12.0 Hz, 2H), 3.10 (t, *J* = 6.0 Hz, 1H), 2.66 (s, 3H), 2-63-2.54 (m, 2H), 2.34 (s, 3H), 2.03-1.92 (m, 3H), 1.65-1.61 (m, 1H); HRMS: *m/z* 478.2562 [M+H]^+^, calcd. for C_30_H_32_N_5_O; Found 478.2602.

#### Compound 1q

To a degassed solution of intermediate **5a** (1 equiv.) and 4**-**formylbenzene boronic acid (1 equiv.), and 2M aq. K_2_CO_3_ solution (1.5 mL/mmol) in 1,4-dioxane (4.5 mL/mmol) in a microwave vial was added Pd(PPh_3_)_4_ (0.05 equiv.) and the mixture was heated at 100°C for 2 min. using microwave. The reaction mixture was diluted using water, extracted using CH_2_Cl_2_, and the combined organic layers concentrated under reduced pressure and purified over Silica gel to afford **1q**. ^1^H NMR (400 MHz, CDCl_3_): δ 9.18 (s, 1H), 8.92 (s, 1H), 8.56 (s, 1H), 8.51 (d, *J* = 5.04 Hz, 1H), 8.08 (d, *J* = 8.00 Hz, 1H), 7.63 (d, *J* = 7.88 Hz, 2H), 7.48 (d, *J* = 7.88 Hz, 2H), 7.25-7.21 (m, 4H), 7.09-7.05 (m, 1H), 3.49 (s, 2H), 2.37 (s, 3H), 2.32 (s, 6H); HRMS: *m/z* 396.2144 [M+H]^+^, calcd. for C_25_H_26_N_5_; found 396.2197.

#### Compound 1r

Prepared by Suzuki reaction of **5a** with **7** as described above for **1q**. ^1^H NMR (400 MHz, CDCl_3_) of **1r**: d 9.14 (s, 1H), 8.92 (s, 1H), 8.52 (d, *J* = 4.52 Hz, 2H), 8.13 (d, *J* = 6.0 Hz, 1H), 7.60 (d, *J* = 8.56 Hz, 2H), 7.48 (d, *J* = 8.56 Hz, 1H), 7.25-7.21 (m, 2H), 7.10-6.96 (m, 3H), 3.34-3.23 (m, 8H), 1H), 2.81 (s, 3H), 2.34 (s, 3H), 2.10-1.64 (m, 4H); HRMS: *m/z* 437.2409 [M+H]^+^, calcd. for C_27_H_29_N_6_; found 437.2449.

#### Compound 1s

Prepared by amide formation between **5c** and **6j** using EDC/HOBt coupling. ^1^H NMR (600 MHz, CDCl_3_): d 8.49 (d, *J* = 6.0 Hz, 1H), 8.38 (s, 1H), 8.15 (d, *J* = 6.0 Hz, 1H), 7.99 (br s, 1H), 7.88-7.83 (m, 4H), 7.68 (d, *J* = 6.0 Hz, 1H), 7.53 (t, *J* = 6.0 Hz, 1H), 7.42-7.38 (m, 2H), 7.33 (d, *J* = 12.0 Hz, 1H), 7.26 (d, *J* = 6.0 Hz, 1H), 7.21 (d, *J* = 6.0 Hz, 1H), 7.08 (t, *J* = 6.0 Hz, 1H), 6.97 (s, 1H), 2.29 (q, *J* = 6.0 Hz, 2H), 2.61 (q, *J* = 6.0 Hz, 2H), 2.40 (s, 3H), 2.38 (s, 6H); HRMS: *m/z* 439.2202 [M+H]^+^, calcd. for C_28_H_30_N_5_O; found 439.2256. Purity (HPLC): >98%.

#### Compound 1t

Prepared by amide formation between **5c** and **6k** using EDC/HOBt coupling. MS of **1t**: *m/z* 464.23 [M+H]^+^, calcd. for C_28_H_29_N_6_O; found 465.24.

### 2.2. Synthesis of IMT isomer 2 and analogs 2a-b

Prepared using intermediates **5a**-**c** (Scheme 3A).

#### IMT isomer 2

Buchwald coupling of intermediate **8** with amine **9** as described above for **1** afforded intermediate **10**. MS of **10**: *m/z* 409.15 [M]^+^, calcd. for C_24_H_19_N_5_O_2_; found 410.16 [M+H]^+^.

Intermediate **10** underwent reductive amination with 4-methylpiperazine using NaCNBH_3_ as described for **1o** to give IMT isomer **2**. ^1^H NMR (400 MHz, CDCl_3_+ CD_3_OD) of **2**: δ 9.15 (s, 1H), 8.81 (s, 1H), 8.46 (d, *J* = 4.88 Hz, 1H), 8.35 (s, 1H), 8.27 (d, *J* = 6.88 Hz, 1H), 8.07 (d, *J* = 7.64 Hz, 2H), 7.85 (m, 1H), 7.57 (d, *J* = 8.0 Hz, 1H), 7.47 (d, *J* = 8.8 Hz, 1H), 7.45 (d, *J* = 8.0 Hz, 2H), 7.37 (m, 1H), 7.24 (d, *J* = 8.0 Hz, 1H), 7.15 (d, *J* = 4.96 Hz, 1H), 3.58 (s, 2H), 2.52 (br s, 8H), 2.33 (s, 6H); MS: *m/z* 494.2624 [M+H]^+^, calcd. for C_29_H_32_N_7_O; found: 494.2678.

#### Compounds 2a and 2b

Aldehyde **8** underwent reductive amination with neopentyl and cyclohexyl amine, NaCNBH_3_, and AcOH, as described above for intermediate **10**, and the resulting products were reacted with Boc-anhydride to afford intermediates **11a** and **11b**. Next, Buchwald coupling of **11a** and **11b** with amine **9** as described above for **1** afforded the Boc-protected derivatives of compounds **2a** and **2b**. Finally, Boc-deprotection in the latter products using 4M HCl in dioxane gave the title products **2a** and **2b**. MS of **2a**: *m/z* 481.27 [M+H]^+^. MS of **2b**: *m/z* 493.27 [M+H]^+^.

### 2.2. Synthesis of IMT isomer 3

Prepared using intermediates **12** and **14** as outlined in Scheme 3B.

#### Intermediate 12

2-Aminopyrimidine underwent Buchwald coupling with 3-bromo-4-methylbenzaldehyde to afford intermediate **12**. MS of **12**: *m/z* 213.09 [M]^+^, calcd. for C_12_H_11_N_3_O; found 214.09 [M+H]^+^.

#### Intermediate 14

To a solution of 3-aminopyridine (1 equiv.) and DIEA (3 equiv.) in dry THF (5 mL/mmol) was added 4-chloromethylbenzoyl chloride (1.2 equiv.) at room temperature and the resulting mixture was stirred for 16 hours to afford intermediate **13** after purification. MS of **13**: *m/z* 246.06/248.05 [M]^+^, calcd. for C_12_H_11_ClN_2_O; found 247.06/249.06 [M+H]^+^.

A solution of intermediate **13** (1 equiv.), N-Boc-piperidine (1 equiv.), and DIEA (3 equiv.) in dry THF (5 mL/mmol) was heated at 90 °C for 16 h. Reaction mixture was worked-up using water and CH_2_Cl_2_, and the combined organic layers concentrated under reduced pressure and chromatographed over Silica gel using CH_2_Cl_2_-MeOH-aq. NH_3_ to afford Boc-protected-**14**. The latter underwent Boc deprotection to afford **14**. MS: *m/z* 296.16 [M]^+^, calcd. for C_17_H_20_N_4_O; found 297.17 [M+H]^+^.

#### IMT isomer 3

A solution of **12** and **14** in methanol underwent reductive amination using NaCNBH_3_ and AcOH to give IMT isomer **3**. ^1^H NMR (400 MHz, CDCl_3_+ CD_3_OD) of **3**: d 8.68 (s, 1H), 8.37 (s, 1H, and d, *J* = 4.6 Hz, 2H), 8.30 (s, 2H), 7.83 (d, *J* = 7.13 Hz, 2H), 7.82 (s, 1H), 7.43 (d, *J* = 7.88 Hz, 2H), 7.31 (dd, *J* = 7.96, 4.6 Hz, 1H), 7.17 (d, *J* = 7.64 Hz, 1H), 7.01 (d, *J* = 7.48 Hz, 1H), 6.89 (s, 1H), 6.69 (t, *J* = 4.72 Hz, 1H), 3.57 (s, 4H), 3.53 (s, 4H), 2.29 (s, 3H); HRMS: *m/z* 494.2624 [M+H]^+^, calcd. for C_29_H_32_N_7_O; found: 494.2672.

### 2.2. Screening and evaluation of IMT isomers and analogs(Sun et al. 2019; Sinha et al. 2019)

N2a695 cells were used to screen all new compounds and in the follow-up studies with compounds found active in the preliminary screen. In a typical experiment, 6-well tissue culture plates (Corning) were seeded at 4.0×10^5^ – 4.5×10^5^ N2a695 cells/mL, 2 mL/well for overnight incubation. When cells were >95% confluent, media were exchanged with fresh media containing 10 µM solutions of compounds and cells were incubated at 37 °C in 5% CO_2_ for 5 hours. Culture media were collected and soluble Aβ concentrations in the media were determined by ELISA or MSD plates for human Aβ Peptides as per manufacturer instructions. Signals for Aβ were measured using Perkin Elmer Envision and SQ120 MSD ELISA reader. Follow-up studies with N2a695 cells were performed similarly.

### 2.3. Effects of IMT and isomers on APP metabolism(Netzer et al. 2017; Sun et al. 2019; Sinha et al. 2019)

N2a695 cells were treated with compounds for 5 hours as described above, and media were aspirated out (or collected for determination of Aβ levels). Cells were scraped in cold Dulbecco’s PBS buffer (1 mL) containing mini EDTA-free protease inhibitor (Roche) and centrifuged for 1 minute at 13,000 rpm at 4 °C to form a cell pellet. The buffer was aspirated and the cell pellets were lysed in 3% SDS plus protease inhibitor cocktail by sonication on ice for two rounds of 20 seconds on a low setting. Protein concentrations were measured using the Pierce BCA Protein Assay (Thermo Fisher) kit in accordance with the manufacturer’s instructions.

To perform WBs, N2a695 cell lysates from 1a and analogs-treated samples were run on a 10-20% or a 16.5 % Tris-Tricine gel (Criterion) and electro transferred to PVDF membranes (EMD Millipore) overnight at 30V. PVDF membranes were incubated in PBS containing 0.25% glutaraldehyde (Sigma) for 30 min after electro transference, blocked for 30 minutes in milk PBST, incubated with primary antibody RU369 for 1 hour at room temperature followed by washing and incubation with an HRP-linked secondary antibody and detected with enhanced chemiluminescence ECL reagents. WB images were analyzed using ImageJ to quantify the prominent bands.

To determine effects of compounds on BACE1 vs. GS inhibition, we used N2a cells transiently transfected with full length APP (APP-FL) or with APP99 (APP-βCTF) as described previously(Netzer et al. 2017; Sun et al. 2019). After 48 hours, media were removed and fresh media containing compound 1a and analogs were added. Following 5 hours of incubation, cell supernatants were collected, and analyzed using MSD-ELISA for Aβ and for sAPPα and β.

### 2.4. In vivo brain permeability and retention of IMT analogs(Sun et al. 2019)

All procedures involving animals were approved by The Rockefeller University Institutional Animal Care and Use Committee and were in accordance with the National Institutes of Health guidelines. Mesylate salts of the isomeric IMT analogs (1 or 3 mg/mL in water, 125 µL, 50 mg/kg) were administered intraperitoneally (i.p.) or through oral gavage to 8 weeks old C57BL/6J WT mice. Mice were euthanized 4 hours post drug administration and brain hemispheres and plasma were harvested and collected in pre-weighted tubes and snap-frozen in liquid nitrogen. To measure brain and plasma concentrations of the specific compounds, mouse brain tissue was homogenized and extracted using Ethanol, and plasma samples were extracted using Acetonitrile. Concentration of the drug and metabolites in brain and in plasma was determined by LC-MS/MS analysis.

### 2.5. Drug Extraction from Brain(Sun et al. 2019)

After tubes were weighed to calculate brain weight and thawed to room temperature, 1 mL of EtOH (200 Proof) was added to the microcentrifuge tubes containing the harvested right brain hemispheres. 10 µL of 1 µM internal standard (ABG190, a synthetic analog of 1a) was added to each tube and samples were sonicated to homogeneity (∼2 min). Tubes were shaken at 40 min at room temperature (1K RPM) and centrifuged for 8 minutes at 13K RPM. The supernatant (0.9 mL) was transferred to a new collection tube and 0.5 mL EtOH was added to the pellet for a second round of extraction as described above. 600 µL of the supernatant was combined with the first collection before samples were submitted for LCMS-MS analysis.

### 2.6. Drug Extraction from Blood(Sun et al. 2019)

300 µL of acetonitrile was added to collected blood samples. 10 µL of 1 µM internal standard (ABG190) was added to each tube and samples were sonicated to homogeneity (∼2 min). Tubes were contributed at 13K RPM for 9 minutes. 300 µL of the supernatant was collected and combined with 500 µL of 5 mM ammonium formate before samples were submitted for LCMS-MS analysis.

### 2.7. In vitro kinase activity assay(Jester et al. 2010)

The assay was performed by Luceome Biotechnologies, LLC. Typically, 10 mM stock solutions of the compounds were diluted in DMSO to a concentration of 250 μM. Prior to initiating the assay, all test compounds were evaluated for false positive against split-luciferase(Jester et al. 2010). For kinase assays, each Cfluc-Kinase was translated along with Fos-Nfluc using a cell-free system (rabbit reticulocyte lysate) at 30 °C for 90 min. 24 μL aliquot of this lysate containing either 1 μL of DMSO (for no-inhibitor control) or compound solution in DMSO (10 μM final concentration) was incubated for 30 minutes at room temperature followed by 1 hour in presence of a kinase specific probe. 80 μL of luciferin assay reagent was added to each solution and luminescence was immediately measured on a luminometer. The percent Inhibition was calculated using the following equation: % Inhibition = (ALUcontrol– ALUsamplex 100)/ALUcontrol.

## 3 Results

### 3.1 Chemistry

IMT isomers **1**-**3** possess all five rings and the chemical functions that broadly match the parent compound (Fig. 1). We designed these isomers by making one or two hypothetical fragmentations across C-N and C-C bonds and re-joining the resulting fragments through other ring(s) and keeping the functionalities similar to IMT, as outlined in Scheme 1. Arrow ‘a-c’ shown in RED and the double arrow shown in BLUE denote the site(s) of fragmentation and re-attachment of various bonds, respectively. Thus, the left part (three ‘ABC’ rings) of IMT will separate from the right two ‘DE’ rings involving a C-N bond fragmentation (designated by ‘a’, Eqn. 1) next to ring ‘C’, and the resulting left fragment, **I**, will connect through ring ‘A’ (pyridine ring) to amide ‘N’ of the right fragment, **II**, giving isomer **1**. Alternatively, IMT will undergo two cleavages across the C-C bond between ring A and B (step ‘b’) for both isomers **2** and **3**, and another (1) C-C bond cleavage between -C(O)- and ring ‘D’ (step ‘c’, Eqn. 2) for isomer **2** and (2) C-N bond cleavage next to ring ‘C’ (step ‘a’, Eqn. 3) for isomer **3**. These cleavages would give three fragments each, **III**, **IV** and **V** for isomer **2**, and **III**, **VI** and **II** for isomer **3**. Note that **II** is a common fragment for both the isomers **1** and **3**, and **III** is common for isomers **2** and **3**. To generate isomer **2**, fragment **III** will combine with the amide carbon in **IV** and the ring ‘B’ of fragment **IV** with ring ‘D’ of **V**, both involving the C-C bond formation (Eqn. 2). Finally, fragment **III** will combine (C-N bond formation) with the amide nitrogen in **II**, and the ring ‘C’ of fragment **VI** with ‘N-Me’ ring ‘E’ of **II** (C-C bond formation) giving isomer **3** (Eqn. 3).

**Scheme 1.**
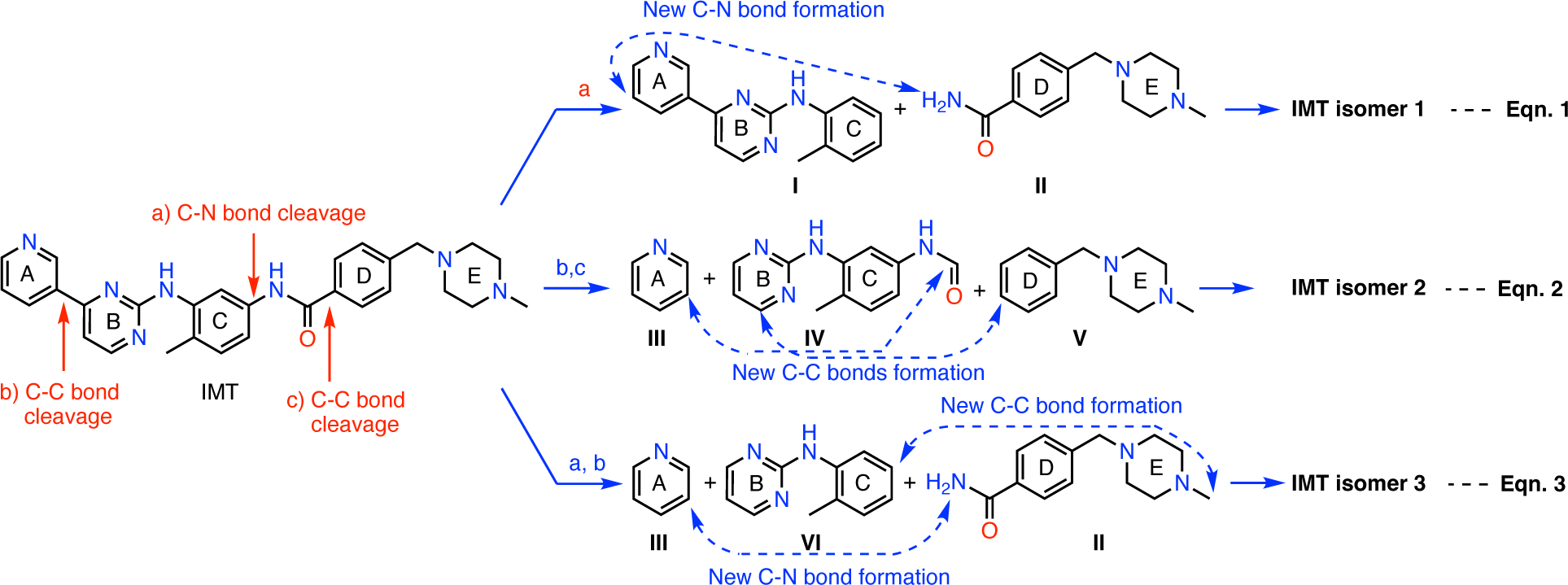
Design of IMT isomers **1**, **2** and **3**. Shown are hypothetical fragmentation of IMT involving (a) C-N bond or (b, c) C-C bond cleavage giving fragments **I**-**VI**, and re-assembly of these fragments to afford IMT isomers **1**, **2** and **3**. Note: fragment **II** is common for both IMT isomers **1** and **3**, and **III** for isomers **2** and **3**. Key: Red arrow, site of C-C or C-N bond cleavage for fragmentation; Blue arrow, C-C or C-N bond connection for re-assembly of the molecules.

Synthesis of IMT isomers **1**-**3** and analogs. We prepared IMT isomer **1** and its analogs **1a**-**1t** using the readily available intermediates, as outlined in Schemes 2A-D. First, to prepare IMT isomer **1**, intermediate **4a** was reacted with o-toluidine and the resulting product **5a** underwent Buchwald coupling(Ruiz-Castillo and Buchwald 2016) with amide **6** to give isomer **1** (Scheme 1A). Similarly, intermediate **4b** reacted with o-toluidine to give **5b**, and both **5a** and **5b** underwent Buchwald coupling(Ruiz-Castillo and Buchwald 2016) with various amides **6a**-**i** giving products **1a**-**n**, several after Boc deprotection as needed (Scheme 2A and Table 1). Analogs **1o** and **1p** were prepared by reaction of **1l** and **1m** with formaldehyde under the reductive amination conditions using NaCNBH_3_ (Scheme 2B). Analog **1q** was prepared by Suzuki coupling of **5a** with 4-formylphenylboronic acid, **7a**, followed by reductive amination of the resulting product with dimethylamine, and **1r** by Suzuki coupling of **5a** with boronic acid **7b** (Scheme 2C)(Miyaura and Suzuki 1995). The remaining analogs of **1**, i.e., **1s** and **1t** were obtained by reacting **4c** with 3-aminopyridine, followed by Boc-deprotection giving amine **5c** and reacting the latter with acids **6j** and **6k** (Scheme 2D).

**Scheme 2.**
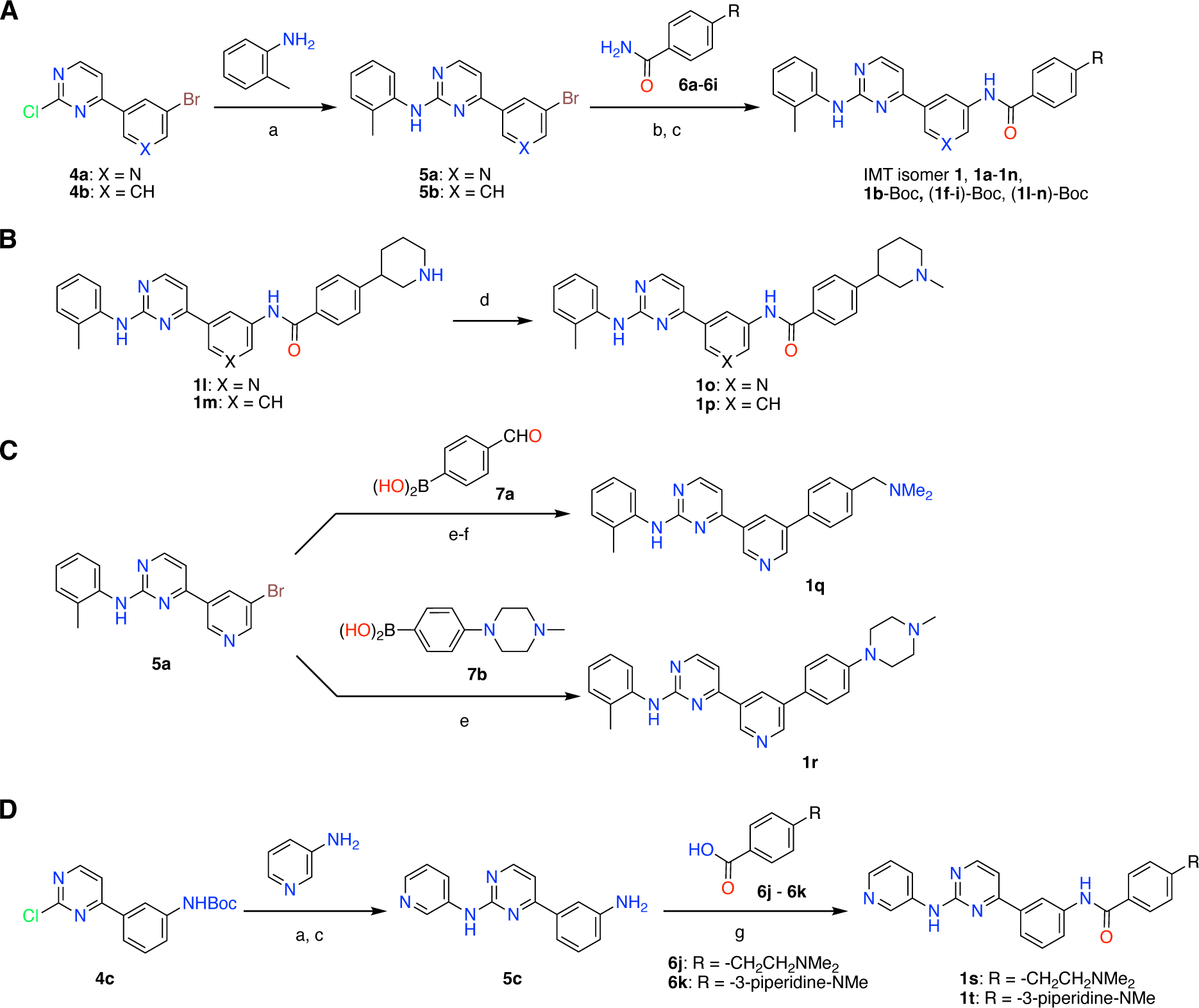
Synthesis of IMT isomer 1 and analogs 1a-t. Key: a) 3N HCl, Dioxane. microwave, 100°C, 2 h. b) Pd_2_(dba)_3_, XanthPhos, Cs_2_CO_3_, 1,4-Dioxane, microwave, 100°C. c) 4M HCl in dioxane, EtOAc, RT, 2 h. d) CH_2_O, NaCNBH_3_, DCE, 0 °C - RT, 16 h. e) Pd(PPh_3_)_4_, aq. K_2_CO_3_, 1,4-Dioxane, microwave, 100°C, 2 h. f) Me_2_NH, Na(OAc)_2_BH, AcOH, DCE. g) EDC, HOBt, CH_2_Cl_2_, RT, 16 h. All compounds possess original A-E lettering for IMT shown in Fig. 1 and Scheme 1.

Next, we prepared IMT isomer **2** and its analogs **2a**-**b** using intermediates **8** and **9**, as described in Scheme 3A. Intermediates **8** and **9** reacted together under the Buchwald coupling conditions affording **10**, which underwent reductive amination with N-methylpiperazine to give IMT isomer **2**. Alternatively, intermediate **8** underwent reductive amination with cyclohexyl amine and neopentyl amine and Boc-protection of the resulting amines to give intermediates **11a** and **11b**, which reacted with intermediate **9** under the Buchwald coupling conditions, followed by Boc-deprotection to give analogs **2a**-**b**. Finally, to prepare IMT isomer **3**, we prepared intermediate **12** by reacting 3-bromo-4-methylbenzaldehyde with 2-amino-pyrimidine under Buchwald conditions, and intermediate **14** by reacting 4-chloro-mthylbenzoyl chloride with 3-aminopyridine 3-amino-pyridine and then with N-Boc-piperazine, followed by N-deprotection. Subsequently, we coupled intermediates **12** and **14** together under the reductive amination conditions using NaCNBH_4_ to give the title product **3**(Afanasyev et al. 2019).

**Scheme 3.**
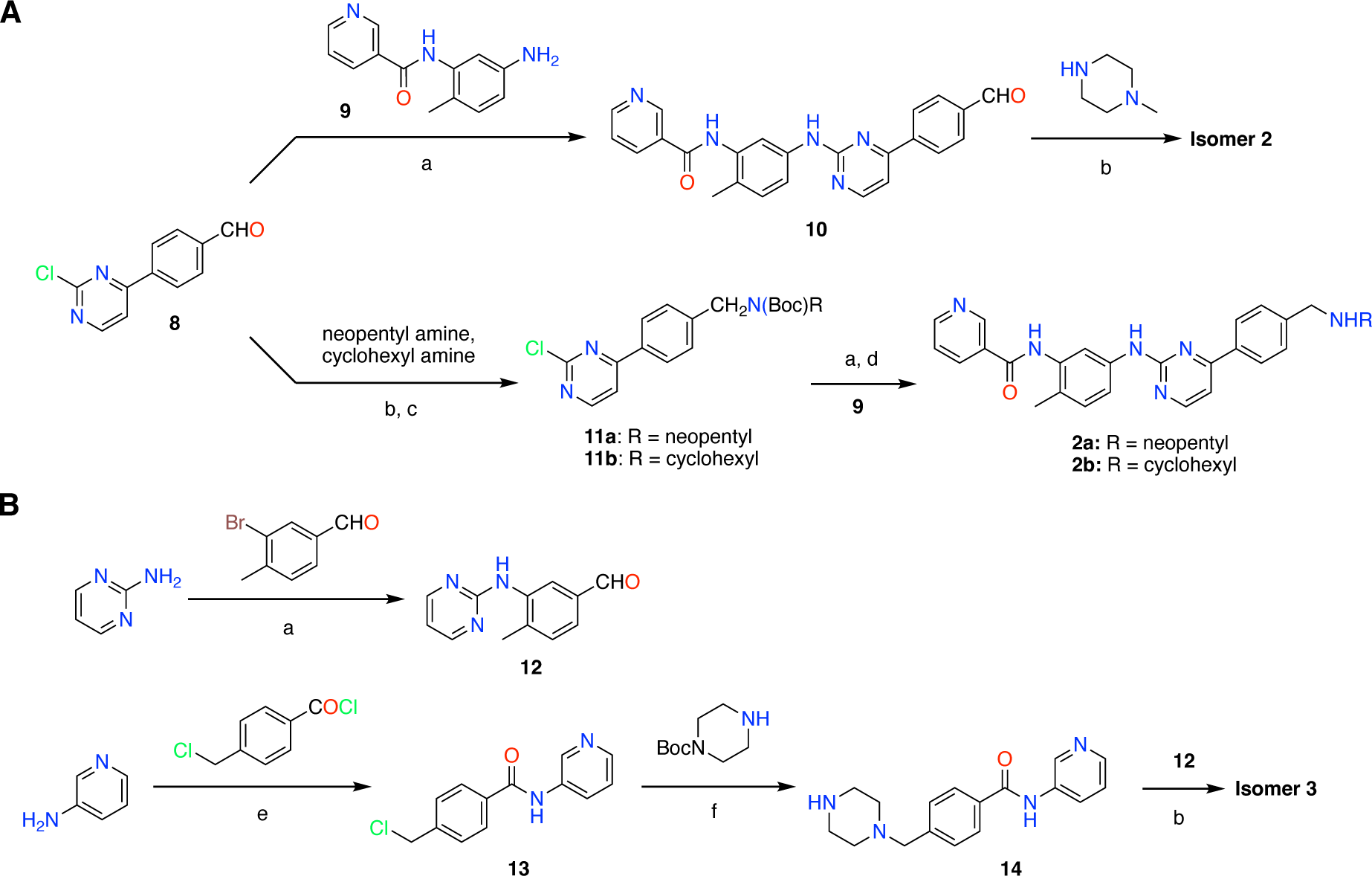
Synthesis of IMT isomers **2** and **3**. Key: a) Pd(dba)_3_, XanthPhos, Cs_2_CO_3_, 1,4-Dioxane, microwave, 100°C, 2 h. b) Na(OAc)_3_BH, DCE, AcOH. c) Boc_2_O, ACN. d) 4M HCl in dioxane, EtOAc, RT, 2 h. d) Methanolic HCl, EtOAc. e) DIEA, THF, RT, 3 h. f) DIEA, THF, 90°C, 2 h.

### 3.2 Structural diversity

The majority of IMT isomer **1** analogs, including **1a**-**1n** and both analogs of isomer **2**, i.e., **2a** and **2b**, differ from one-another in ring ‘A’ and/or in ‘E’ (See: Fig. 1 for lettering of the rings, which are based on the original A-E lettering for IMT shown in Scheme 1). New IMT isomer **1** analogs contain piperazine ring, a cyclic amine or piperidine ring connected through C-C or C-N bond to ring D, while all other isomer **1** and both isomer **2** analogs possess a substituted alkylamine instead of the ring E. These modifications were made to compare the similar changes made in IMT analogs. There was no additional difference between two analogs, **2a** and **2b**, of isomer **2**. All 20 analogs of isomer **1** also have rings ‘A-D’ and their arrangement is similar with three exceptions. 1) Nine compounds possess 1,3-substituted benzene and the remaining 11 analogs contain 3,5-substituted pyridine (Py) as the middle ring ‘A’, 2) The first ring from the left (ring ‘C’) in 2 analogs, **1s** and **1t**, is 3-aminopyridine instead of o-toluidine in all remaining 18 compounds. 3) Analogs **1q** and **1r** do not possess the ‘amide group’ that connects the middle ring ‘A’ to the 4th ring ‘D’.

### 3.3 Evaluation

Previously, we showed that two chemically distinct compounds, IMT and DV2-103 lower Aβ production primarily by reducing BACE processing of APP(Netzer et al. 2017). Similarly, numerous analogs of IMT also lowered Aβ production by reducing BACE processing of APP(Sun et al. 2019). In the present study we further examined the effects of these compounds on γ-secretase catalyzed Aβ formation and compared these to the more radically isomeric analogs of IMT.

Our results show that IMT, DV2-103, and IMT isomer **1** are γ-secretase modulators; i.e. These compounds favor production or inhibition of different lengths of Aβ peptides (differing in their C-termini). Specifically, we exposed N2a695 cells to increasing concentrations of each compound and measured the production of Aβ38, 40, and 42. IMT, DV2-103 and IMT isomer **1** (Fig. 2A) inhibit the formation of Aβ38 least, compared to Aβ40 and 42, and even boost levels of Aβ38 above controls at a drug concentration of 5μM. Remarkably, this occurs for all Aβ peptides tested shorter than 40 amino acids (Fig. 2B,C)). Thus, both IMT, DV2-103, and IMT isomer **1** are γ-secretase modulators. Remarkably, while each compound modulates γ-secretase activity, most conspicuously at 5μM, this effect vanishes at 10μM, relative to controls.

**Fig. 2.**
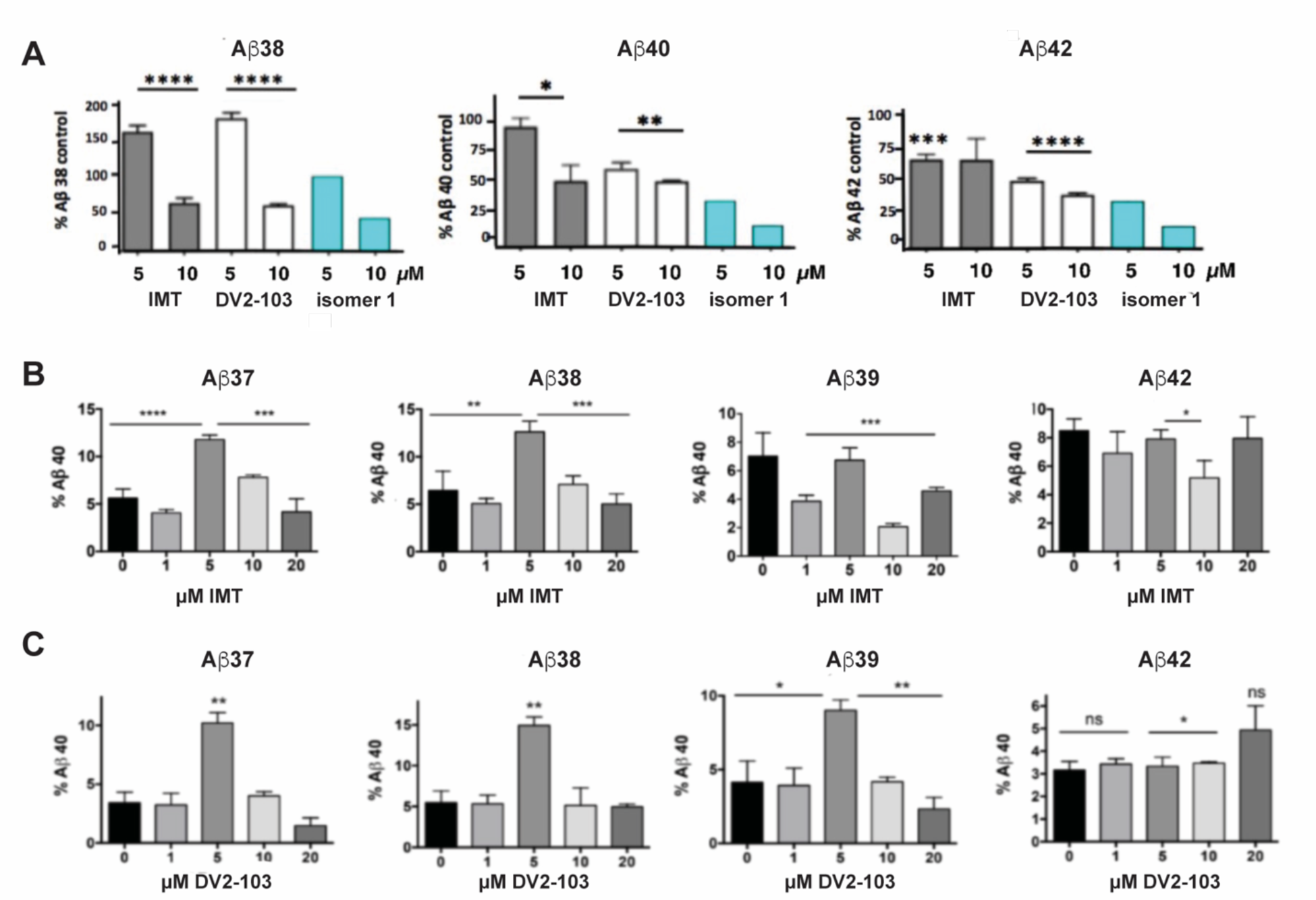
IMT, DV2-103, and IMT isomer **1** are γ-secretase modulators. (A) N2a695 cells incubated with IMT, DV2-103 and IMT isomer 1 lower levels of Aβ40 and 42 more than Aβ38, especially at a drug concentration of 5μM, as measured by ELISA. Means differ significantly for IMT and DV2-103 compared to DMSO controls, N = 3 x 3. Data for Isomer 1 are from a representative sample. Differences between means for (A)are analyzed by One-way Anova for IMT and DV2-103 treated cells. Differences among means comparing 5μM IMT and DMSO controls (B,C)) are analyzed by Student’s T test (S.E.M.).

Subsequently, we screened all IMT derivatives shown in Table 1 for levels of Aβ40, Aβ38, and Aβ42 peptides. We found that the majority of compounds reduced production of Aβ40 and Aβ42 more than Aβ38 peptide (Table 2 and Supporting Information (SI) Fig. S-1). This indicates that isomer **1** and analogs modulate γ-secretase cleavage of C-terminal APP since the differences in lengths of these peptides is determined by γ-secretase according to differences in utilization of the APP γ-secretase cleavage sites.

**Table 2.** Isomeric IMT analogs are γ-secretase modulators^a^.

| Comp. ID | % Aβ38, 40, 42 of DMSO ctrl at 10 (5 μM) conc. | Comp. ID | % Aβ38, 40, 42 of DMSO ctrl at 10 (5 μM) conc. | Comp. ID | % Aβ38, 40, 42 of DMSO ctrl at 10 (5 μM) conc. |
| --- | --- | --- | --- | --- | --- |
| **1a** | 86, 40, 35 (105, 57, 49) | **1b** | 61, 34, 33 (95, 56, 53) | **1c** | 91, 46, 44 (99, 57, 54) |
| **1d** | 80, 29, 26 (113, 52, 45) | **1f** | 80, 43, 43 (106, 64, 58) | **1g** | 85, 41, 46 (106, 56, 58) |
| **1h** | 123, 73, 65 (112, 83, 75) | **1i** | 92, 38, 38 (120, 60, 56) | **1k** | 22, 14, 11 (60, 34, 28) |
| **1l** | 52, 21, 17 (107, 48, 43) | **1o** | 95, 42, 38 (112, 63, 56) | **1s** | 24, 18, 15 (58, 41, 38) |
| **1t** | 47, 23, 23 (89, 45, 42) | **2b** | 74, 71, 74 (93, 90, 88) |  |  |

<sup>a</sup> The screening experiment was performed for Aβ40, 38 and 42 in cell supernatants of N2a695 cells using MSD ELISA. Drug concentrations are 10 μM or 5 μM. Aβ values are expressed as percentages of control Aβ38, 40, and 42, respectively. Results for the IMT isomer 1 derivatives shown here were obtained from a single experiment.

To test whether these compounds lower Aβ levels by affecting the BACE and/or γ-secretase cleavages, we examined their effects using wild-type (wt) cells transfected with APP-FL and APP βCTF (C99), respectively. BACE inhibitor, MK8931, and γ-secretase inhibitor, DAPT, were used as controls and the experiment was performed and processed as described previously^13^. The results shown in Fig. 3A and 3B revealed that all 4 compounds reduced β- and γ-cleavages of APP similarly to IMT. There were reductions in Aβ production in both cases, but more so in cells transfected with full-length APP indicating that these compounds, like IMT (Netzer et al. 2017), reduce both BACE and γ-secretase cleavages of APP but that attenuation of BACE processing accounted for the greater part of Aβ reduction (Fig. 3A, B). None of these compounds showed any toxicity to N2a695 cells at 10µM concentration (Fig. 3C). This was assessed by measuring the percentage of viable cells in the drug treated groups compared to the DMSO control after 5 hours incubation under the conditions used for the Aβ assay.

**Fig. 3.**
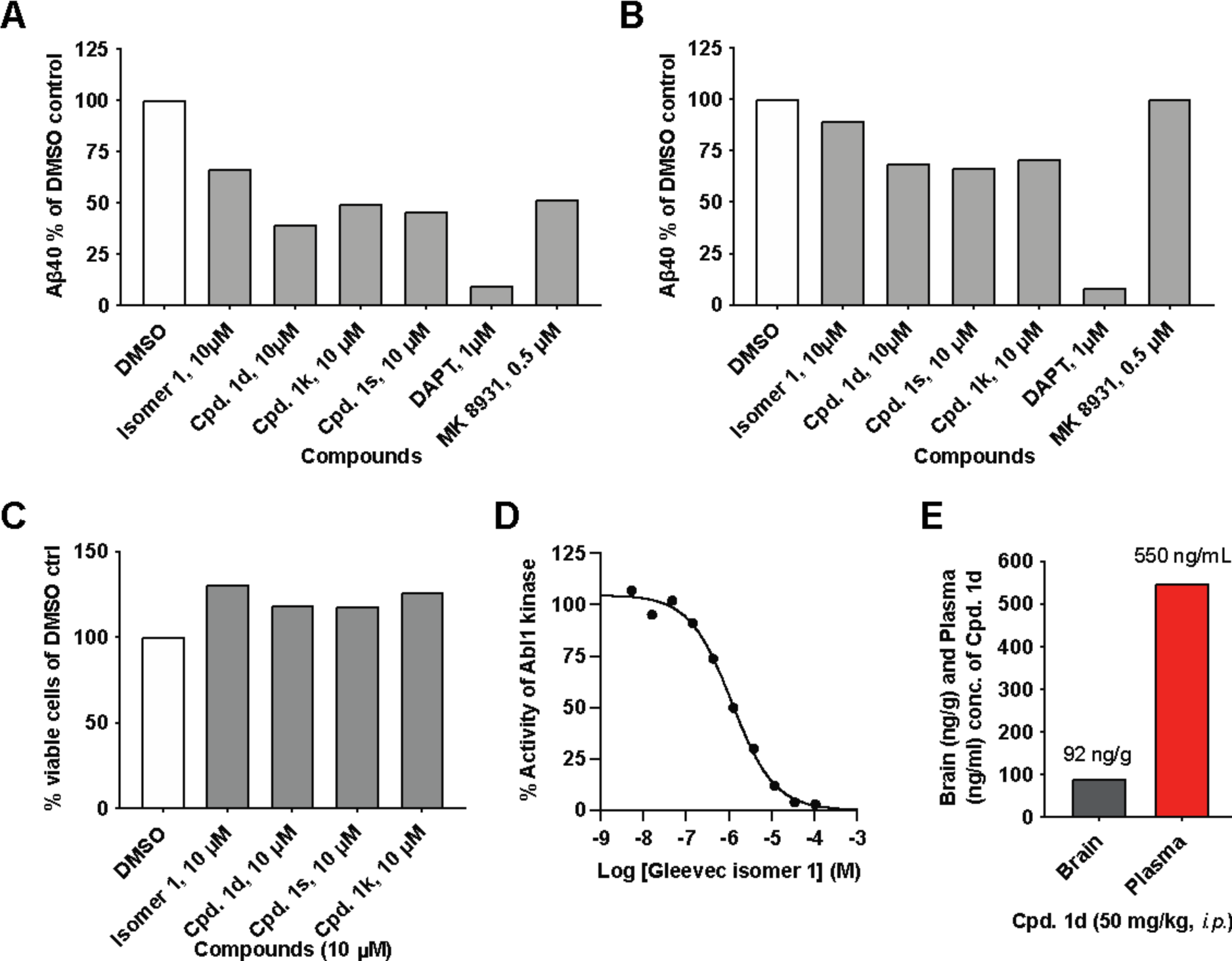
IMT isomer **1** and its analogs **1d**, **1k** and **1s** lower BACE and γ-secretase cleavage of APP and lower levels of Aβ in N2a cells transiently transfected with APP 695 (left graph) or APP C99 (right graph): (A) full length APP695 (APP-FL) or (B) APP C99 (β-CTF). (C) Percentage of viable wild-type (WT) N2a cells upon treatment with IMT isomers **1**, cpd. **1d**, cpd. **1k** and cpd**. 1s** compared to DMSO control under the same conditions used to test Aβ production. (D) Effects of isomer **1** on Abl1 kinase in vitro. (E) Brain and plasma concentrations of isomer **1** and cpd. **1d** in 2 months old WT mice 4 hours after i.p. injection of 50 mg/kg of each drug. Data for A-C are from representative samples.

IMT inhibits Abl1 kinase with low nanomolar affinity. Earlier, we prepared and evaluated numerous IMT analogs to find that many of these analogs reduced Aβ levels in cells similarly to IMT, while inhibiting Abl kinase less potently, compared to IMT. In other words, there isn’t a good correlation between the Abl kinase inhibitory activity vs. the Aβ lowering effects in cells contacted with the IMT analogs. We have evaluated IMT isomer **1** to find that it inhibits Abl kinase less potently (IC_50_: 1.172 µM) (Fig. 3D) than IMT IC_50_: 0.038 µM)(Buchdunger et al. 1996), while it reduced Aβ levels more potently than IMT (Fig. 1C). This result further reinforces our prior observation that there is no or little connection between the Aβ-lowering activity of IMT and its inhibition of Abl1 kinase(Netzer et al. 2003). Finally, we tested the brain permeability of compound **1d** by administering it to 2 months old mice. Plasma and brain tissue were collected 4 hours post drug administration, and LC-MS/MS analysis of the acetonitrile and ethanol extracts was used to measure drug concentration. Compound **1d** possesses similarity to IMT isomer-**1a** and is isomeric to an ABG-179(Sun et al. 2019) analog that possessed a benzene instead of the pyridine (A) ring (see: Fig. 1 for the ring numbering). Earlier, we have shown that ABG-179 possesses superior brain exposure compared to IMT and reduced both Aβ40 and 42 levels significantly in AD mice when delivered acutely for 5 days, and now found that compounds **1d** and ABG-179 possess comparable brain exposure but the latter possess superior plasma half-life (Fig. 3E)(Sun et al. 2019).

## 4 Discussion

Based on the number and variety of chemically distinct compounds that produce the same biochemical effects on APP metabolism(Druker et al. 1996; Nagar et al. 2002; Netzer et al. 2017) and that all active compounds are active at low micromolar concentration, we postulate that IMT, DV2-103 and their analogs are likely to produce their effects on APP metabolism by virtue of their physical rather than stereological properties. For example, physical properties would include acting as a weak base that would cause these molecules to be lysosomotropic. We came to this conclusion by showing that the effects of IMT and DV2-103 on APP metabolism are dependent on acidified lysosomes(Netzer et al. 2017). Reduced dependence on stereological factors would suggest that IMT, DV2-103 and their active derivatives might bind to a polyspecific receptor where binding is less dependent on structural and electrostatic complementarity.

As a first step to test this, we designed derivatives of IMT, referred to here as IMT isomers, and then synthesized derivatives of these compounds and tested their effects on APP metabolism. Our goal was to make a large change in the structure of IMT that would destroy a structural/stereological pharmacophore but still maintain IMT’s physical properties, in particular its property as a weak base, which is necessary for its sequestration in lysosomes through ion trapping(Burger et al. 2015). These properties are preserved in IMT isomer **1**, and we demonstrated that isomer **1** not only lowers levels of Aβ peptide in N2a 695 cells but also lowers BACE processing of APP, and in each case with equal activity or more potently than IMT, while producing the same metabolites of APP in cells exposed to IMT.

Other IMT isomers were synthesized and a subset of these recapitulated IMT’s APP phenotype. Isomers **1** and **1d** also showed increased brain accumulation compared to IMT administered in mice. Also, isomer **1** inhibited Abl kinase activity with over a 100 fold reduction in potency compared to previously published reports of IMT (Buchdunger et al. 1996). Although we had compared the relative effects of γ-secretase and BACE modulation of Aβ generation in cells, we could not rule out that the lowering of Aβ and sAPPβ was not a result of IMT’s effect of stimulating autophagy, since autophagy was previously shown to accelerate lysosomal degradation of APP-βCTF and Aβ(Tian et al. 2011). Nevertheless, we show that IMT, DV2-103, and the IMT isomers tested in this study are modulators of γ-secretase by virtue of the observation that their Aβ-lowering potency differentially affects Aβ peptide lengths depending on drug concentration. Remarkably, Aβ1-42 production is lowered at 5μM drug concentrations, while Aβ1-38 production is inhibited least and in some cases raised. This is important because relative heightened production of Aβ38 has been considered benign, and more recently therapeutic(Cullen et al. 2022), while lowered production of Aβ42 is considered therapeutic; in either case, a decrease in Aβ peptide aggregation may occur.

## 5 Conclusion

In summary, we suggest that IMT and a subset of its isomers, and related lysosomotropic drugs affect APP metabolism through a lysosomal mechanism that increases trafficking of full-length APP directly to lysosomes where it is degraded, thus causing APP to evade processing by BACE and γ-secretase in earlier secretory compartments along the amyloidogenic pathway. The fact that many of these compounds (structurally related or not) are γ-secretase modulators is consistent with a mechanism involving altered trafficking of APP that affects the specificity of γ-secretase cleavage sites in the formation of Aβ peptides.

## 6 Data availability statement

Synthetic methods and analytical data for new compounds and their effects on Abeta levels depicted as a bar graph.

## AUTHOR INFORMATION

Corresponding Authors. William J. Netzer, Anjana Sinha, and Subhash C. Sinha. ORCID 0000-0001-8916-5677

Notes: The authors declare no competing financial interest.

## ACKNOWLEDGEMENTS

We are thankful to Dr. Paul Greengard (Deceased) of the Rockefeller University for his enthusiastic support to this work and Dr. Victor H. Bustos for helpful discussion. Funding support from JPB (#322 and #839 to SCS) and Fisher Center for Alzheimer’s Research Foundation (PG) is duly acknowledged.

