## Supplemental data for "Stretching the structural envelope of isomeric imatinib analogs that reduce β-amyloid production by modulating both β- and γ-secretase cleavages of APP"

**Supporting Information (SI)**

**Contents.**

1. Table 1. HRMS/MS data for Gleevec isomers and analogs S-1
2. HPLC traces for selected compounds S-2
3. Graphical representation of the results from Table 2 (main article) S-10
4. NMR spectra of Gleevec isomers and analogs S-11

**Table S-1. HRMS/MS data for Gleevec isomers and analogs**

| Compound | Molecular Formula | Calcd. m/z [M+H]+ | Found m/z [M+H]+ |
|---|---|---|---|
| **1** | $C_{29}H_{31}N_7O$ | 494.2624 | 494.2653 |
| **1a** | $C_{30}H_{32}N_6O$ | 493.2671 | 493.2716 |
| **1b** | $C_{28}H_{29}N_7O$ | 480.2467 | 480.2506 |
| **1c** | $C_{27}H_{28}N_6O$ | 453.2358 | 453.2403 |
| **1d** | $C_{28}H_{29}N_5O$ | 452.2406 | 452.2450 |
| **1e** | $C_{26}H_{26}N_6O$ | 439.2202 | 439.2243 |
| **1f** | $C_{29}H_{32}N_6O$ | 481.2671 | 481.2716 |
| **1g** | $C_{30}H_{33}N_5O$ | 480.2719 | 480.2776 |
| **1h** | $C_{30}H_{32}N_6O$ | 493.2671 | 493.2718 |
| **1i** | $C_{31}H_{33}N_5O$ | 492.2719 | 492.2772 |
| **1j** | $C_{28}H_{29}N_7O$ | 480.25 | 480.25 |
| **1k** | $C_{28}H_{30}N_6O$ | 479.25 | 479.25 |
| **1l** | $C_{28}H_{28}N_6O$ | 465.2358 | 465.2401 |
| **1m** | $C_{29}H_{28}N_5O$ | 464.2406 | 464.2459 |
| **1n** | $C_{29}H_{29}N_5O$ | 463.24 | 464.24 |
| **1o** | $C_{29}H_{30}N_6O$ | 479.2515 | 479.2560 |
| **1p** | $C_{30}H_{31}N_5O$ | 478.2562 | 478.2602 |
| **1q** | $C_{25}H_{25}N_5$ | 396.2144 | 396.2197 |
| **1r** | $C_{27}H_{28}N_6$ | 437.2409 | 437.2449 |
| **1s** | $C_{28}H_{29}N_5O$ | 439.2202 | 439.2256 |
| **1t** | $C_{28}H_{28}N6O$ | 464.23 | 465.24 |
| **2** | $C_{29}H_{31}N_7O$ | 494.2624 | 494.2678 |
| **2a** | $C_{29}H_{32}N_6O$ | 481.27 | 481.27 |
| **2b** | $C_{30}H_{32}N_6O$ | 493.27 | 493.27 |
| **3** | $C_{29}H_{31}N_7O$ | 494.2624 | 494.2672 |

=====================================================================

Acq. Operator : SYSTEM
Sample Operator : SYSTEM
Acq. Instrument : Prep LC                Location : Pl-C-09
Injection Date : 1/9/2019 1:18:35 PM                Inj : 1
                Inj Volume : 50.000 µl
Acq. Method : D:\ChemStation\l\Methods\Anjana\Reverse and Cyclic Gleevec 010819.M
Last changed : 1/8/2019 2:37:40 PM by SYSTEM
Analysis Method : D:\ChemStation\1\Methods\Emily Mui\Reverse and Cyclic Gleevec 010819.M
Last changed : 1/8/2019 2:38:19 PM by SYSTEM
J
Method Info : Method created 01082019 to run on series of reverse and cyclic gleevec samples at 250 µMin H2o + 0.1% TFA

Sample Info : ABG_95_4 from purified sample        Compound 1

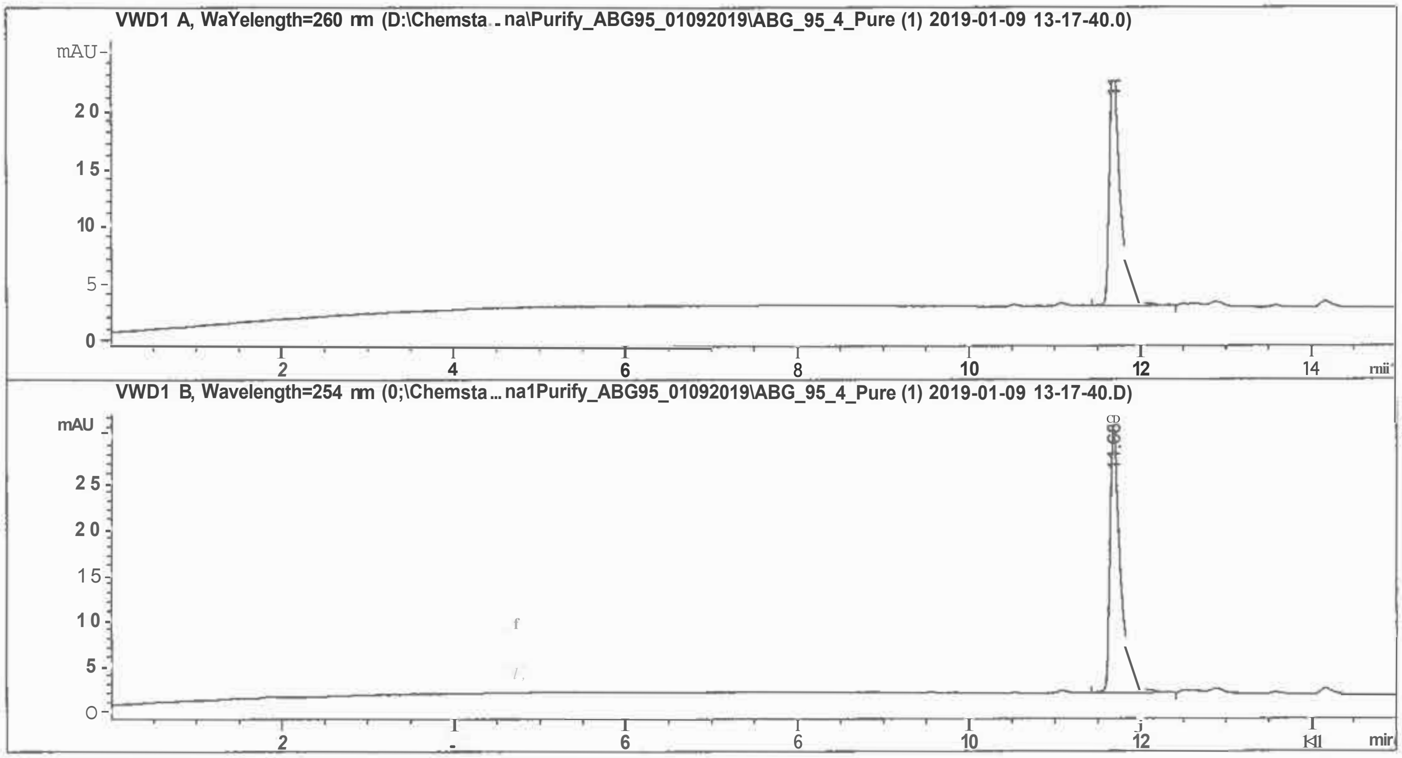


=====================================================================

Fraction Information

=====================================================================

No Fractions found.

=====================================================================

=====================================================================

=====================================================================

Area Percent Report

=====================================================================

Sorted By : Signal
Multiplier : 1.0000
Dilution : 1.0000
Use Multiplier & Dilution Factor with ISTDs

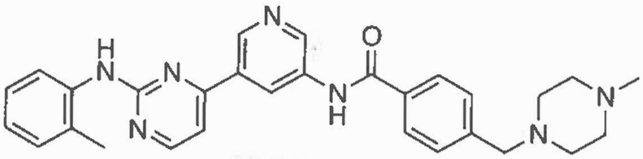

ABG-95

Compound 1

Signal 1: VWDl A, Wavelength=280 nm

| Peak # | RetTime [min] | Type | Width [min] | Area [mAU*s] | Height [mAU] | Area % |
|---|---|---|---|---|---|---|
| 1 | 11.689 | BV R | 0.1240 | 187.49675 | 21.87925 | 100.0000 |

Totals: 187.49675 21.87925

Signal 2: VWDl B, Wavelength=254 nm

| Peak # | RetTime [min] | Type | Width [min] | Area [mAU*s] | Height [mAU] | Area % |
|---|---|---|---|---|---|---|
| 1 | 11.688 | BV R | 0.1239 | 248.65631 | 29.04224 | 100.0000 |

Totals : 248.65631 29.04224

==========================================================================

*** End of Report***

- -Pata File D:\Chemsta ... a\Anjana\Purify_ABG256_1Pure\ABG_256_Pure (1) 2019-01-09 14-42-29.D
sample Name: ABG_256_Pure

Compound 1d

Acq. Operator : SYSTEM
Sample Operator : SYSTEM
Acq. Instrument : Prep LC                Location : Pl-E-08
Injection Date : 1/9/2019 2:43:30 PM     Inj : 1
Inj Volume : 50.000 µl

Method : D:\ChemStation\1\Methods\Anjana\Reverse and Cyclic Gleevec 010819.M
Last changed : 1/8/2019 2:37:40 PM by SYSTEM
Method Info : Method created 01082019 to run on series of reverse and cyclic gleevec samples at 250 µM prepared in H20 + 0.1% TFA.

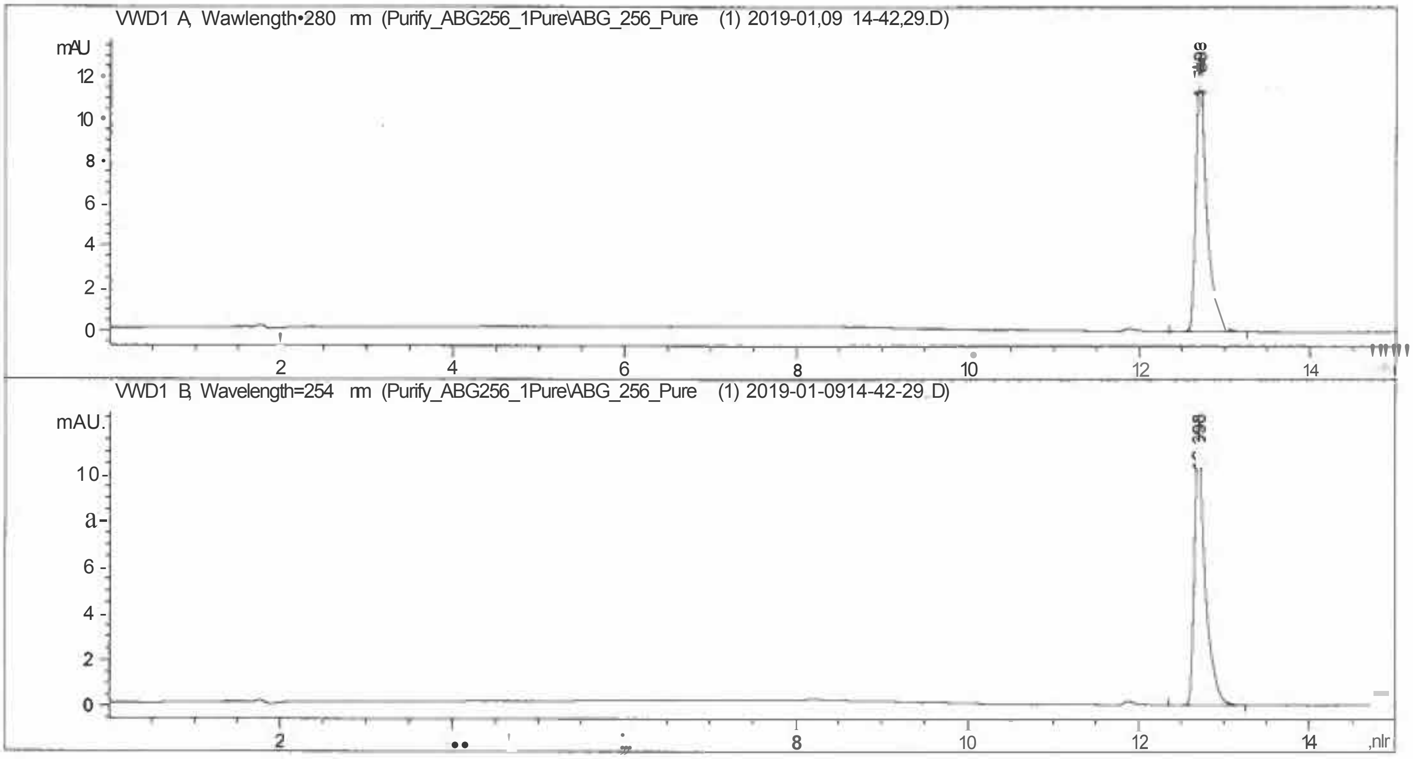


### Fraction Information

No Fractions found.

### Area Percent Report

Sorted By : Signal
Multiplier : 1.0000
Dilution : 1.0000
Use Multiplier & Dilution Factor with ISTDs

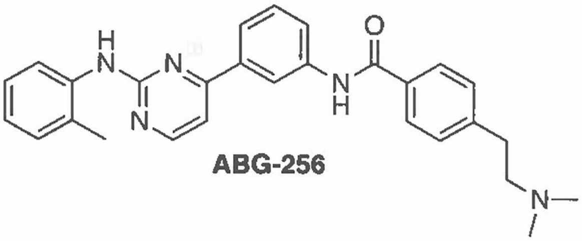


**ABG-256**

Signal 1: VWDl A, wavelength=280 nm

S-4

Compound 1d

| Peak # | RetTime [min] | Type | Width [min] | Area [mAU*s] | Height [mAU] | Area % |
|---|---|---|---|---|---|---|
| 1 | 12,698 | BB | 0.1250 | 113.56995 | 13.11934 | 100.0000 |

Totals : 113.56995 13.11934

Signal 2: VWDl B, Wavelength;254 nm

| Peak # | RetTime [min] | Type | Width [min] | Area [mAU*s] | Height [mAU] | Area % |
|---|---|---|---|---|---|---|
| 1 | 12.698 | BB | 0.1270 | 104.09130 | 12.02237 | 100.0000 |

Totals : 104.09130 12.02237

=====================================================================

*** End of Report***

Sample Name: ABG 260-2-Pure Compound 1m

=====================================================================

Acq. Operator : SYSTEM Seq. Line : 3
Sample Operator : SYSTEM
Acq. Instrument : Prep LC Location : Pl-E-03
Injection Date : 1/9/2019 12:50:12 PM Inj : 1
Inj Volume : 50.000 µl
Different Inj Volume from Sample Entry! Actual Inj Volume : 200.000 µl
Method : D:\ChemStation\l\Data\Anjana\Gleevec\Reinjecting_pure_revandcyc 2019-01-09 12-10-33\Reverse and Cyclic Gleevec 010819.M (Sequence Method)
Last changed : 1/8/2019 2:37:40 PM by SYSTEM
Method Info : Method created 01082019 to run on series of reverse and cyclic gleevec samples at 250 µM prepared in H20 + 0.1% TFA.

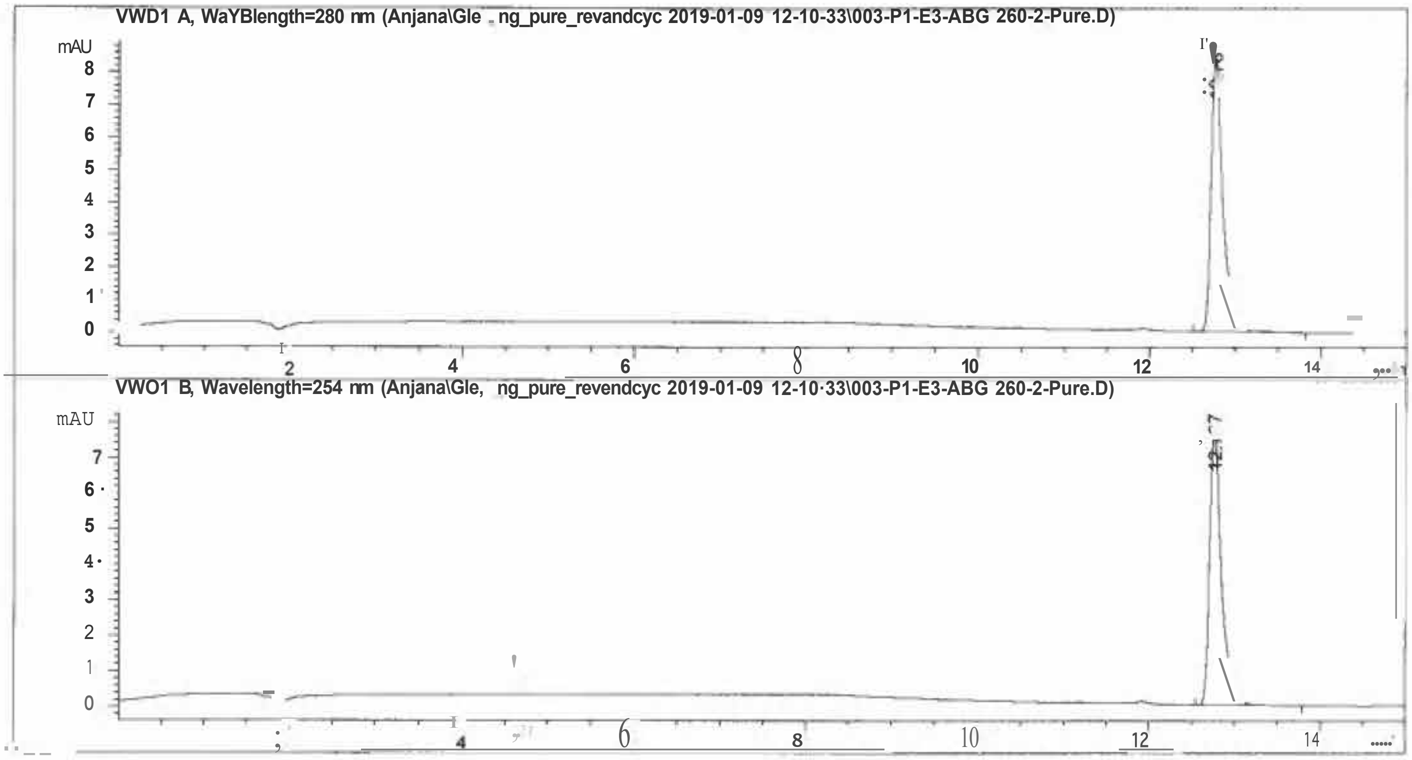


=====================================================================

Fraction Information

=====================================================================

No Fractions found.

=====================================================================

=====================================================================

Area Percent Report

=====================================================================

Sorted By : Signal
Multiplier : 1.0000
Dilution : 1.0000
Use Multiplier & Dilution Factor with ISTDs

Compound 1m

Signal 1: VWD1 A, Wavelength=280 nm

| Peak # | RetTime [min] | Type | Width [min] | Area [mAU*s] | Height [mAU] | Area % |
|---|---|---|---|---|---|---|
| 1 | 12.767 | BB | 0.1387 | 80.65784 | 8.50159 | 100.0000 |
| Totals: | | | | 80.65784 | 8.50159 | |

Signal 2: VWD1 B, Wavelength=254 nm

| Peak # | RetTime [min] | Type | Width [min] | Area [mAU*s] | Height [mAU] | Area % |
|---|---|---|---|---|---|---|
| 1 | 12.767 | BB | 0.1386 | 74.03202 | 7.80763 | 100.0000 |
| Totals: | | | | 74.03202 | 7.80763 | |

*** End of Report***

Data File D:\ChemSta ... nd Cyclic Gleevec 01082019 2019-01-08 14-45-36\011-Pl-Cll-ABG 267.D
Sample Name: ABG 267 Compound 1s

=====================================================================

| | | | |
|---|---|---|---|
| Acq. Operator | : SYSTEM | Seq. Line : | 11 |
| Sample Operator | : SYSTEM | | |
| Acq. Instrument | : Prep LC | Location : | Pl-C-11 |
| Injection Date | : 1/8/2019 5:56:56 PM | Inj : | 1 |
| | | Inj Volume : | 50.000 µl |

Method : D:\ChemStation\1\Data\Anjana\Gleevec\reverse and Cyclic Gleevec 01082019 2019-01-08 14-45-36\Reverse and Cyclic Gleevec 010819.M (Sequence Method)
Last changed : 1/8/2019 2:37:40 PM by SYSTEM
Method Info : Method created 01082019 to run on series of reverse and cyclic gleevec samples at 250 µM prepared in H2O + 0.1% TFA,

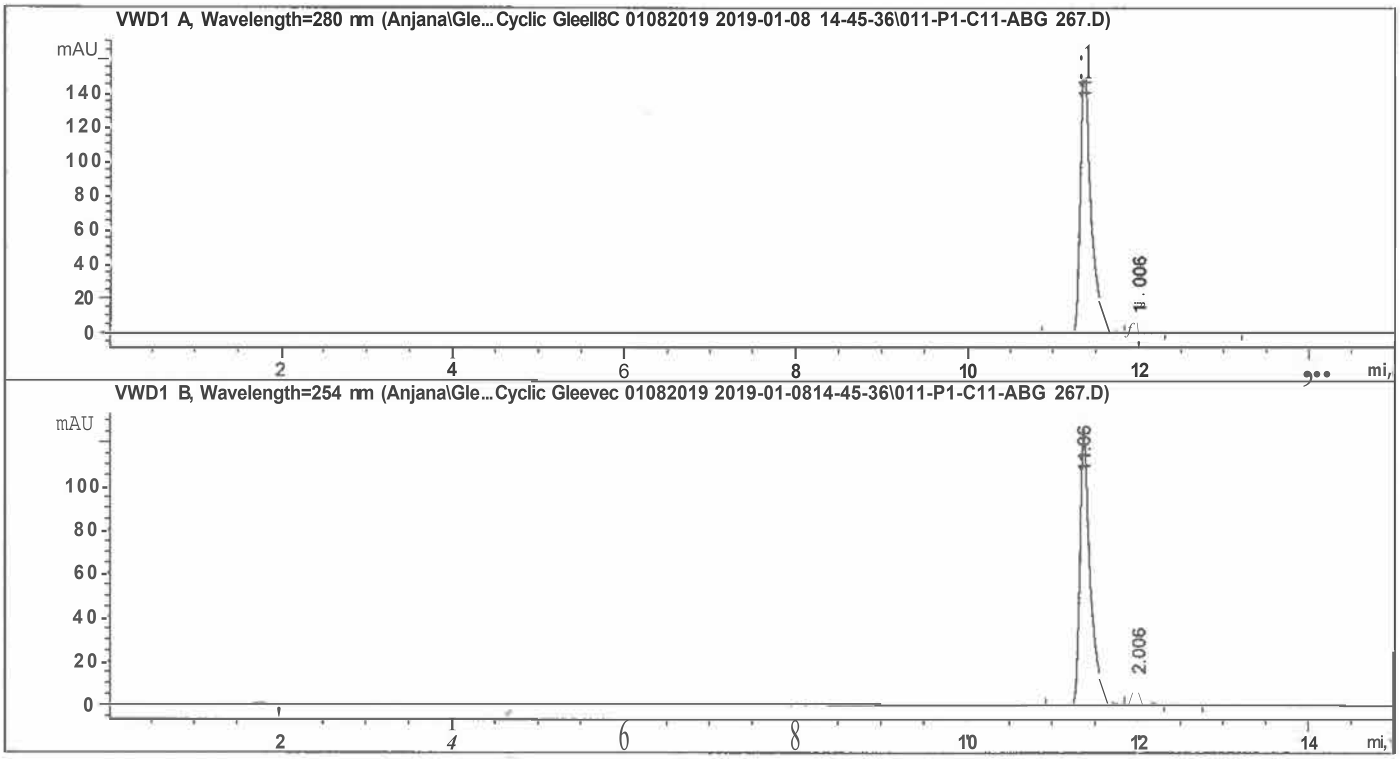


=====================================================================

Fraction Information

=====================================================================

No Fractions found.

=====================================================================

=====================================================================

=====================================================================

Area Percent Report

=====================================================================

| | | |
|---|---|---|
| sorted By | : | Signal |
| Multiplier | : | 1.0000 |
| Dilution | : | 1.0000 |

Use Multiplier & Dilution Factor with ISTDs

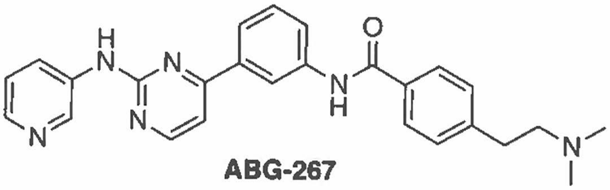

ABG-267

Compound 1s

Signal 1: VWDl A, Wavelength=280 nm

| Peak # | RetTime (min) | Type | Width (min) | Area (mAu•s) | Height (mAU) | Area % |
|---|---|---|---|---|---|---|
| 1 | 11.361 | BV R | 0,1259 | 1418.97791 | 162.51459 | 95.2842 |
| 2 | 12,006 | WE | 0.1269 | 70.22735 | 8,11792 | 4.7158 |

Totals: 1489.20525 170.63251

Signal 2: VWDl B, Wavelength=254 nm

| Peak # | RetTime (min) | Type | Width (min) | Area (mAU*s) | Height (mAU) | Area % |
|---|---|---|---|---|---|---|
| 1 | 11.361 | BV R | 0.1255 | 1101.61719 | 126.61599 | 95.0498 |
| 2 | 12.006 | WE | 0.1246 | 57.37195 | 6.65553 | 4.9502 |

Totals : 1158.98914 133.27152

*** End of Report•••

**Fig. S-1.** Graphical representation of the results from Table 2 (main article).

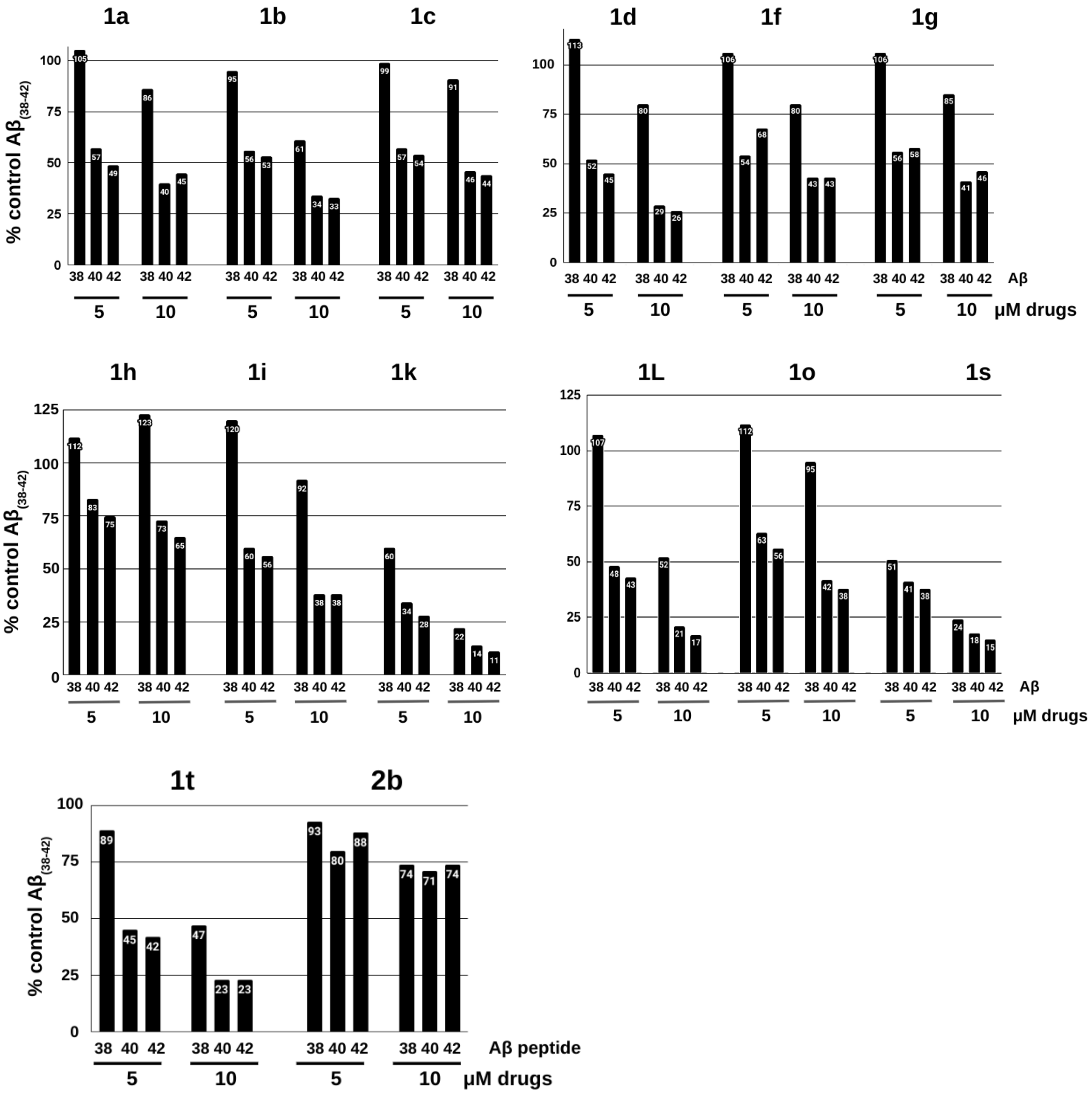


**Note.** Values for compounds' effects on Aβ38, Aβ40, and Aβ42 secreted from N2a695 cells are expressed as percent control (DMSO) for each Aβ peptide.

Type text here

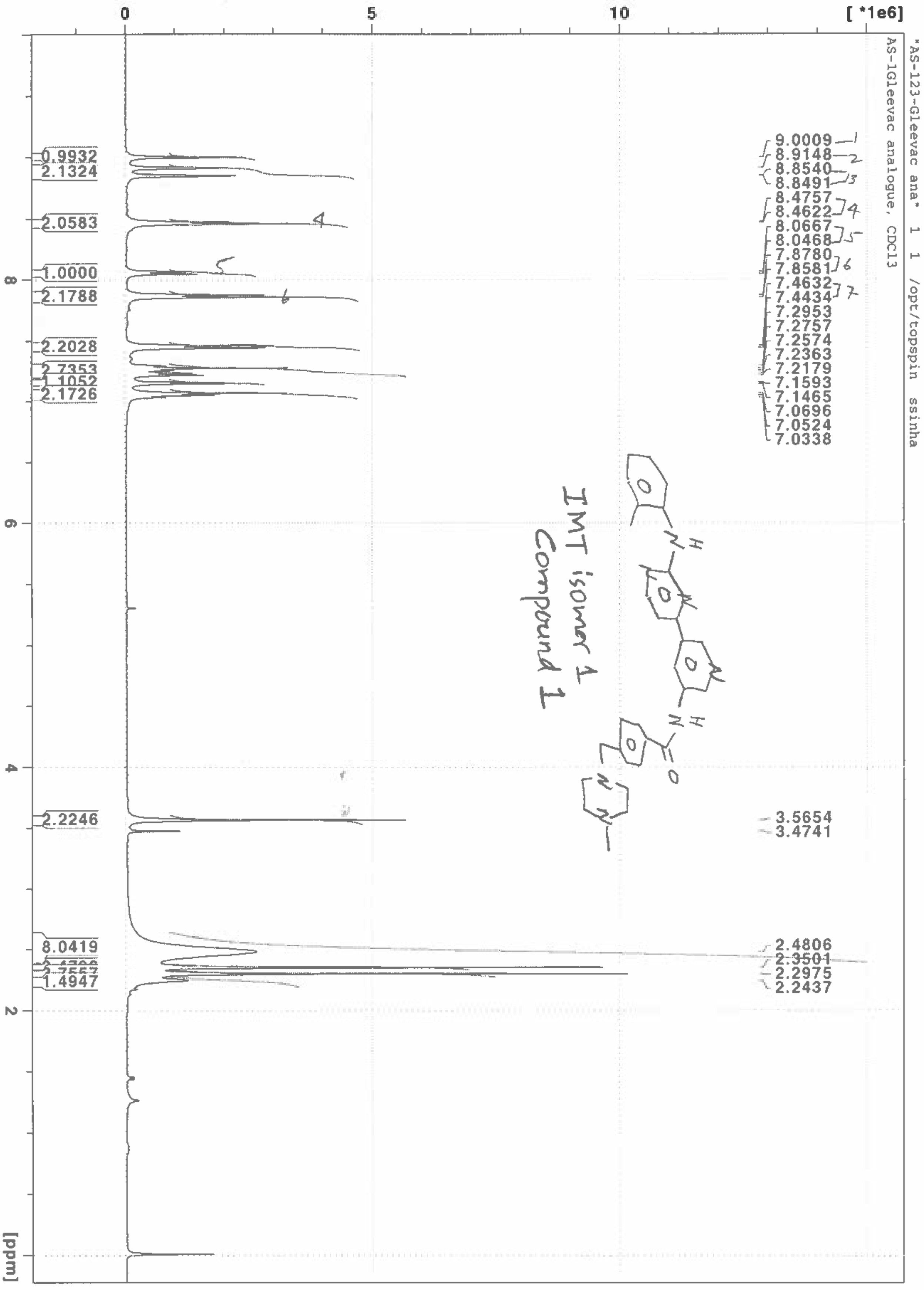
"AS-123-Gleevac ana" 1 1 /opt/topspin ssinha
AS-1Gleevac analogue, CDCl3
IMT isomer 1
Compound 1
9.0009
8.9148
8.8540
8.8491
8.4757
8.4622
8.0667
8.0468
7.8780
7.8581
7.4632
7.4434
7.2953
7.2757
7.2574
7.2363
7.2179
7.1593
7.1465
7.0696
7.0524
7.0338
3.5654
3.4741
2.4806
2.3501
2.2975
2.2437
0.9932
2.1324
2.0583
1.0000
2.1788
2.2028
2.7353
1.1052
2.1726
2.2246
8.0419
1.4947
[ *1e6]
[ppm]

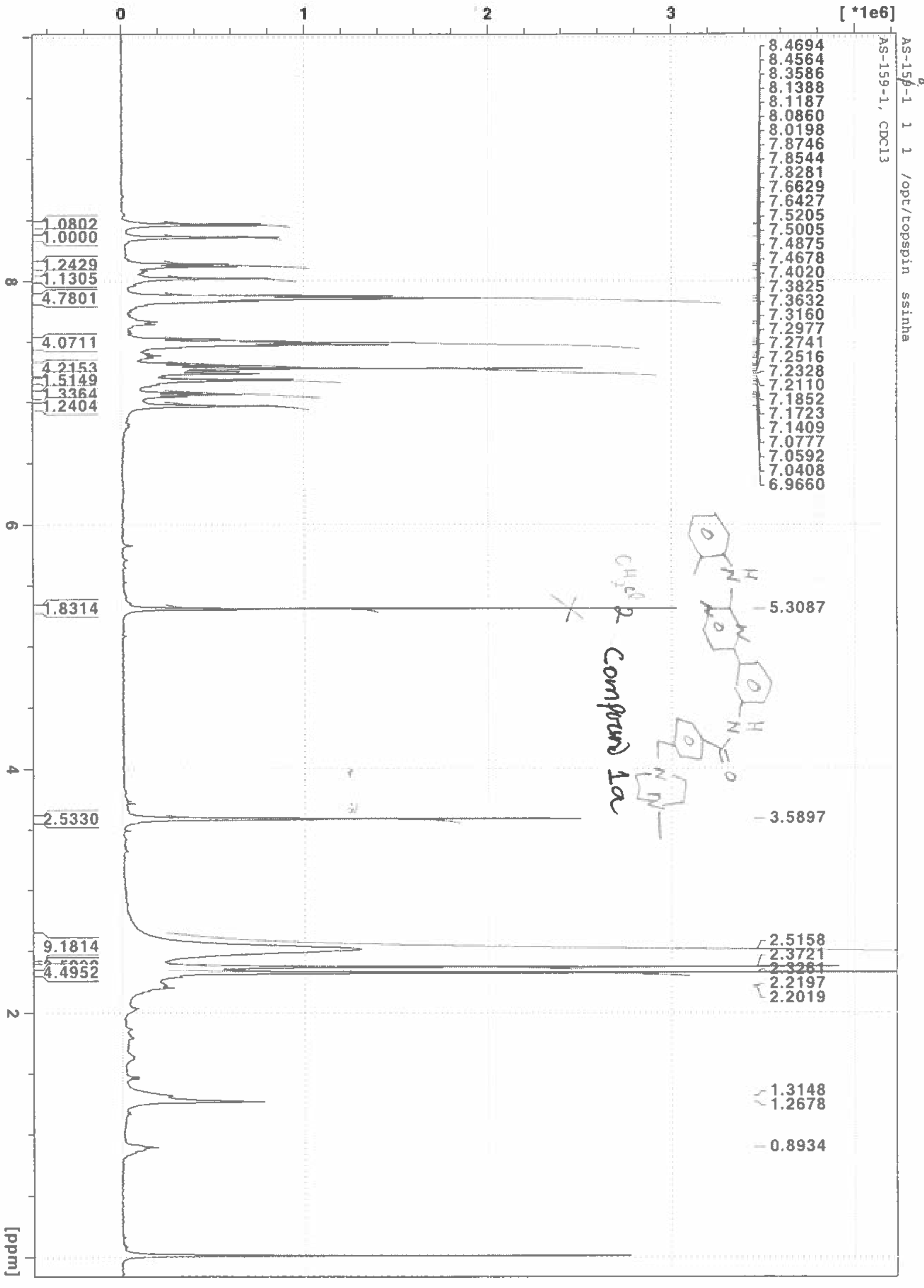
AS-159-1 1 1 /opt/topspin ssinha
AS-159-1, CDCl3
[ *1e6]
[ppm]
8.4694
8.4564
8.3586
8.1388
8.1187
8.0860
8.0198
7.8746
7.8544
7.8281
7.6629
7.6427
7.5205
7.5005
7.4875
7.4678
7.4020
7.3825
7.3632
7.3160
7.2977
7.2741
7.2516
7.2328
7.2110
7.1852
7.1723
7.1409
7.0777
7.0592
7.0408
6.9660
5.3087
3.5897
2.5158
2.3721
2.2197
2.2019
1.3148
1.2678
0.8934
1.0802
1.0000
1.2429
1.1305
4.7801
4.0711
4.2153
1.5149
1.3364
1.2404
1.8314
2.5330
9.1814
4.4952
CH2Cl2
Compound 1a

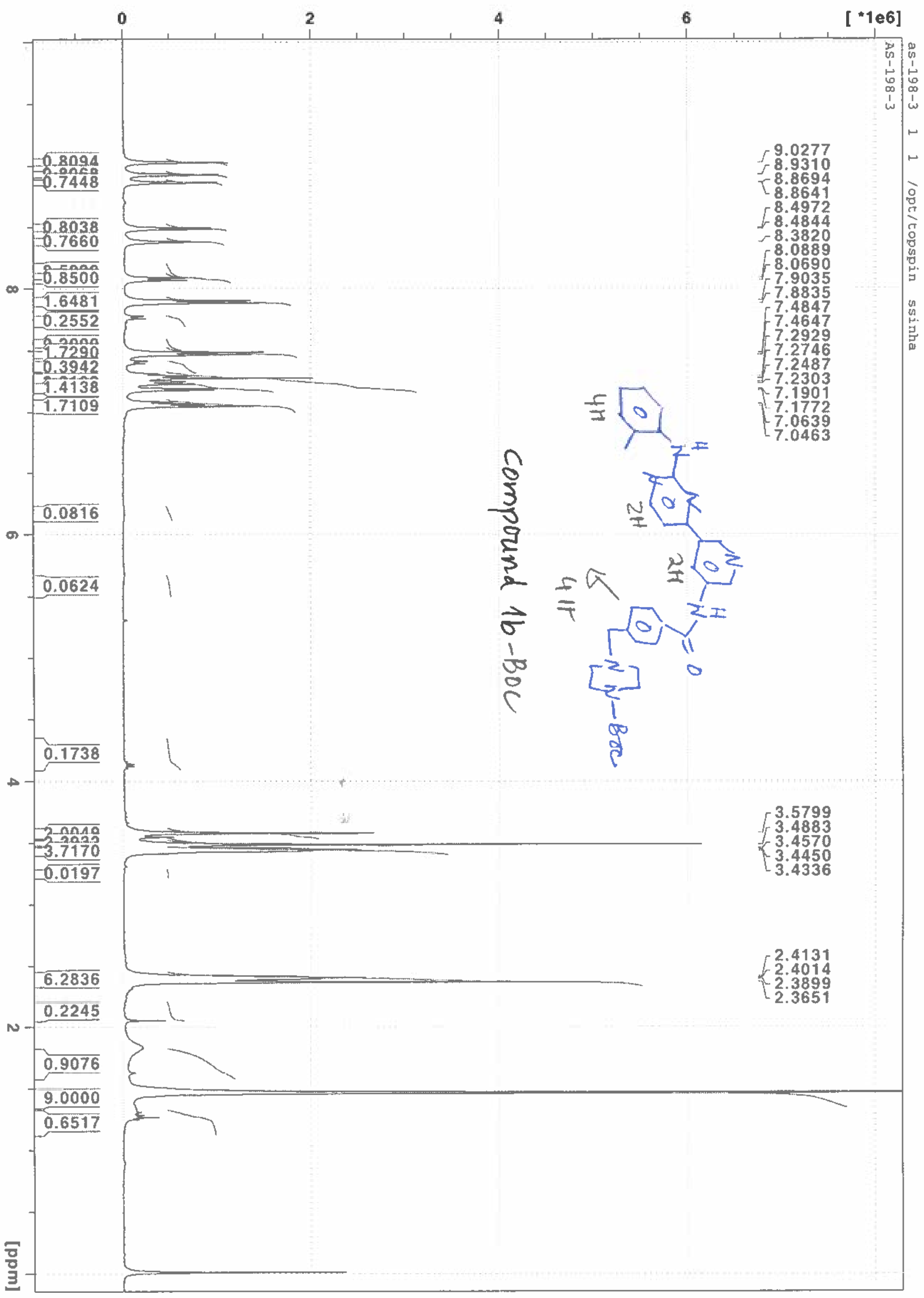
as-198-3 1 1 /opt/topspin ssinha
AS-198-3
[ *1e6]
9.0277
8.9310
8.8694
8.8641
8.4972
8.4844
8.3820
8.0889
8.0690
7.9035
7.8835
7.4847
7.4647
7.2929
7.2746
7.2487
7.2303
7.1901
7.1772
7.0639
7.0463
3.5799
3.4883
3.4570
3.4450
3.4336
2.4131
2.4014
2.3899
2.3651
0.8094
0.7448
0.8038
0.7660
0.8500
1.6481
0.2552
1.7290
0.3942
1.4138
1.7109
0.0816
0.0624
0.1738
3.7170
0.0197
6.2836
0.2245
0.9076
9.0000
0.6517
[ppm]
Compound 1b-Boc



1b-Boc

ABG-255 (01/15/2016)

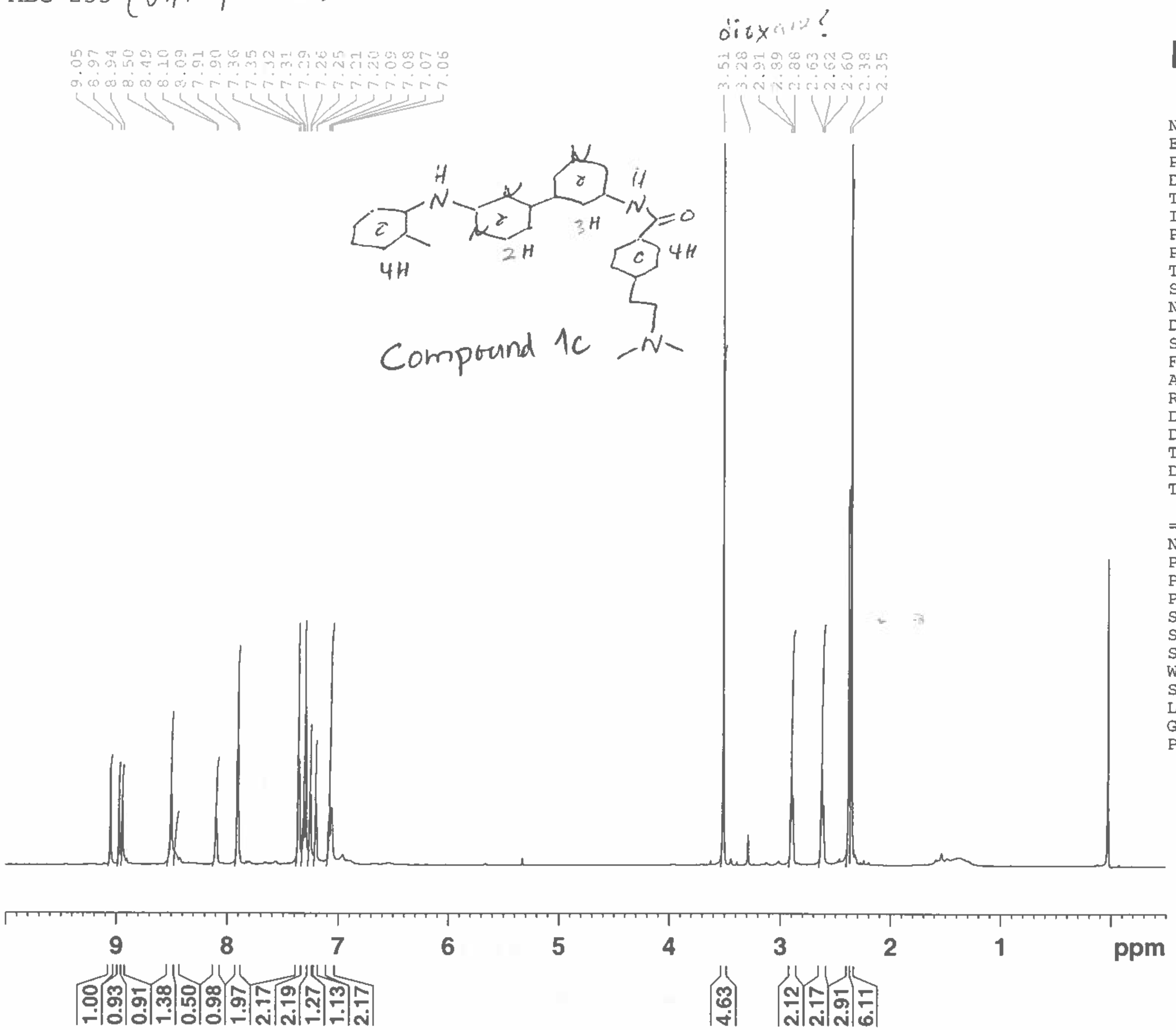


BRUKER

| | |
|---|---|
| NAME | ABG-255 |
| EXPNO | 1 |
| PROCNO | 1 |
| Date_ | 20160121 |
| Time | 16.31 |
| INSTRUM | spect |
| PROBHD | 5 mm CPTCI 1H- |
| PULPROG | zg30 |
| TD | 32768 |
| SOLVENT | CDCl3 |
| NS | 16 |
| DS | 8 |
| SWH | 7788.162 Hz |
| FIDRES | 0.237676 Hz |
| AQ | 2.1038198 sec |
| RG | 9 |
| DW | 64.200 usec |
| DE | 6.00 usec |
| TE | 298.2 K |
| D1 | 1.00000000 sec |
| TD0 | 1 |

======== CHANNEL f1 ========

| | |
|---|---|
| NUC1 | 1H |
| P1 | 7.45 usec |
| PL1 | 4.50 dB |
| PL1W | 5.70400620 W |
| SFO1 | 600.1728538 MHz |
| SI | 16384 |
| SF | 600.1699972 MHz |
| WDW | EM |
| SSB | 0 |
| LB | 1.00 Hz |
| GB | 0 |
| PC | 1.00 |

ABG-256 R (AS-2-85-L) CDCl3

ABG-256

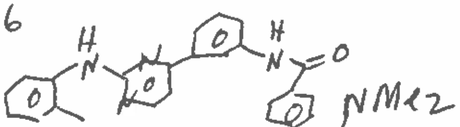

Compound 1d

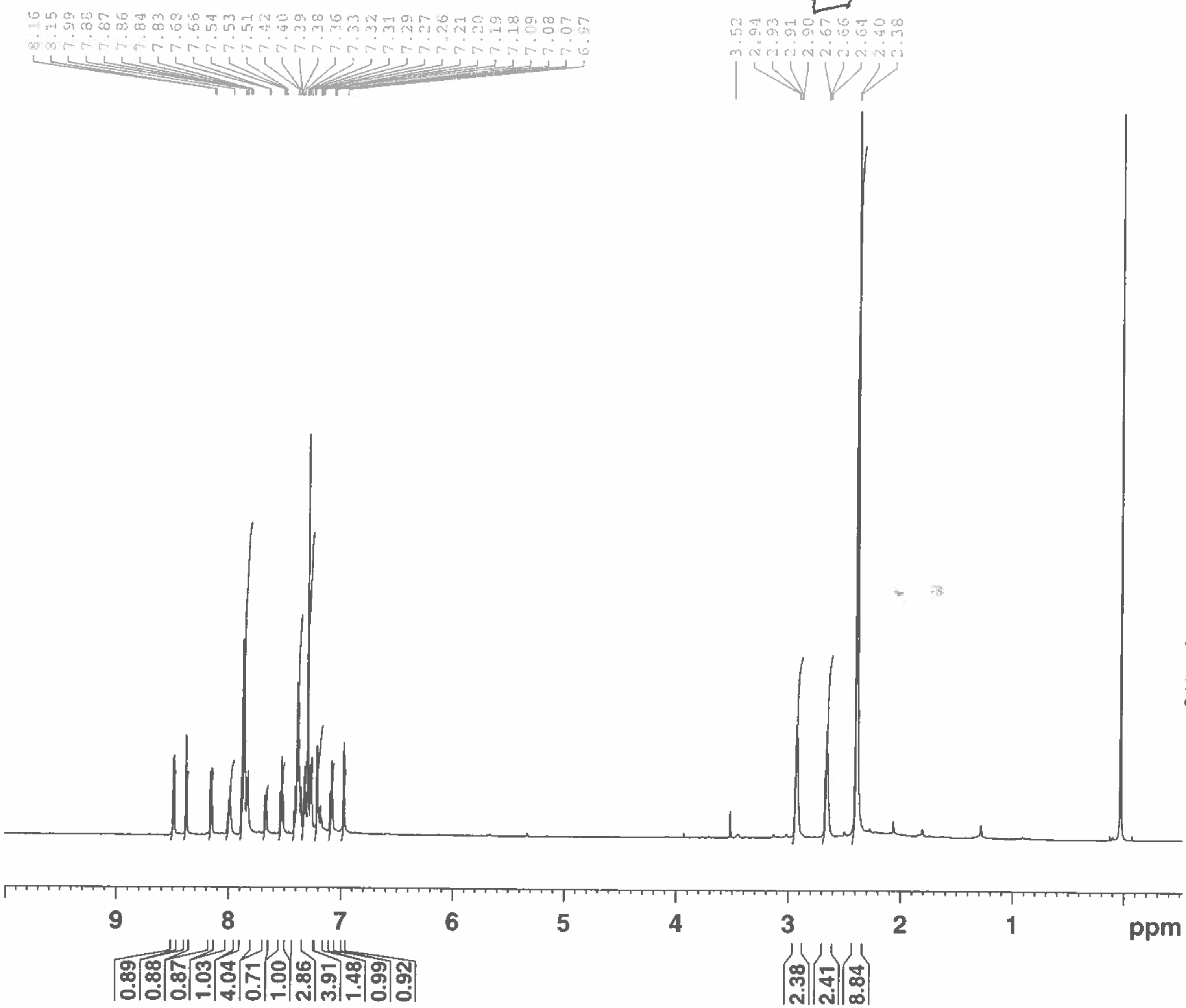


| | |
|---|---|
| NAME | ABG-256R (As2-85L) |
| EXPNO | 1 |
| PROCNO | 1 |
| Date_ | 20160203 |
| Time | 15.26 |
| INSTRUM | spect |
| PROBHD | 5 mm CPTCI 1H- |
| PULPROG | zg30 |
| TD | 32768 |
| SOLVENT | CDCl3 |
| NS | 16 |
| DS | 8 |
| SWH | 7788.162 Hz |
| FIDRES | 0.237676 Hz |
| AQ | 2.1038198 sec |
| RG | 12.7 |
| DW | 64.200 usec |
| DE | 6.00 usec |
| TE | 298.2 K |
| D1 | 1.00000000 sec |
| TD0 | 1 |

======== CHANNEL f1 ========

| | |
|---|---|
| NUC1 | 1H |
| P1 | 7.45 usec |
| PL1 | 4.50 dB |
| PL1W | 5.70400620 W |
| SFO1 | 600.1728538 MHz |
| SI | 16384 |
| SF | 600.1699972 MHz |
| WDW | EM |
| SSB | 0 |
| LB | 1.00 Hz |
| GB | 0 |
| PC | 1.00 |

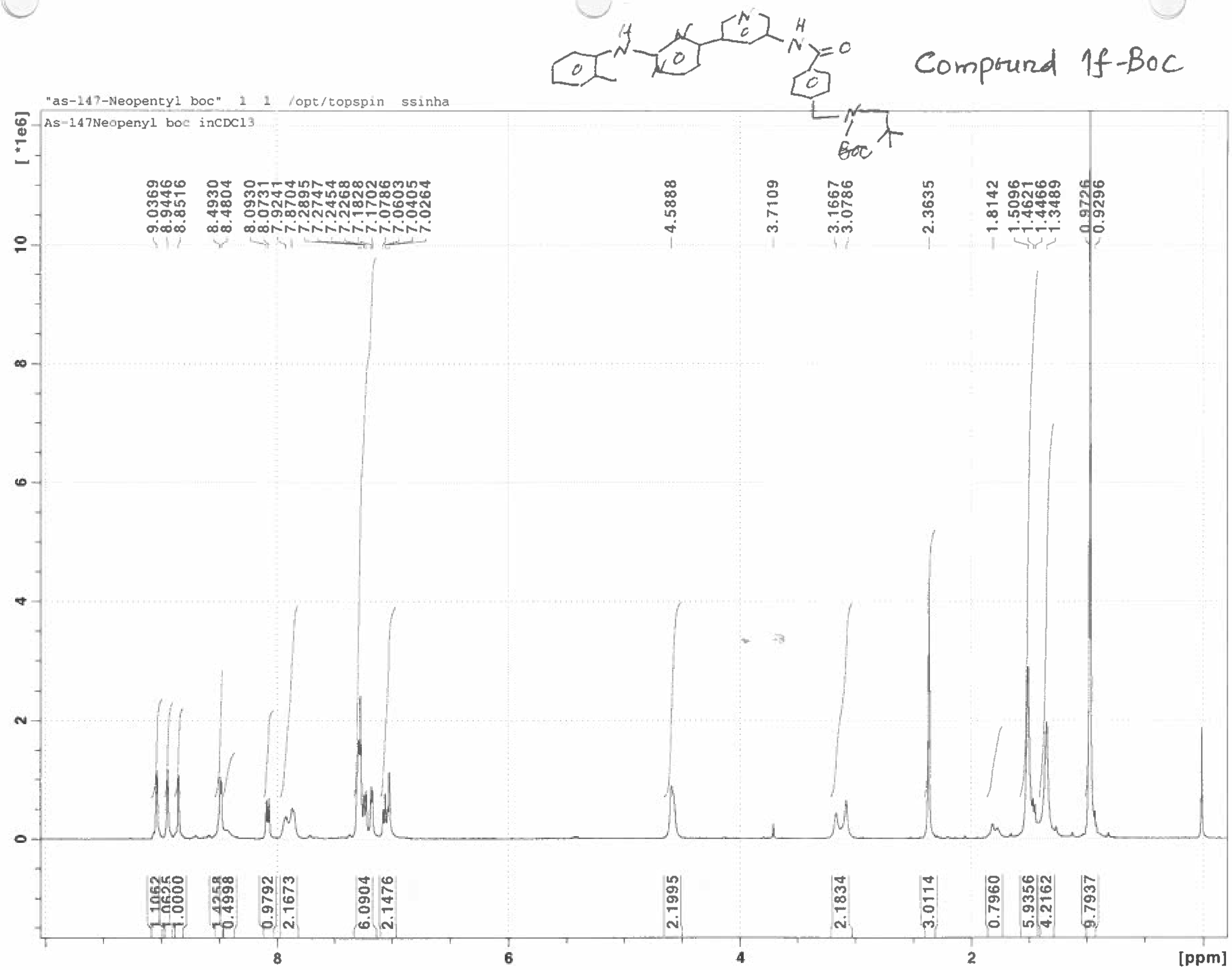


Compound 1f-Boc

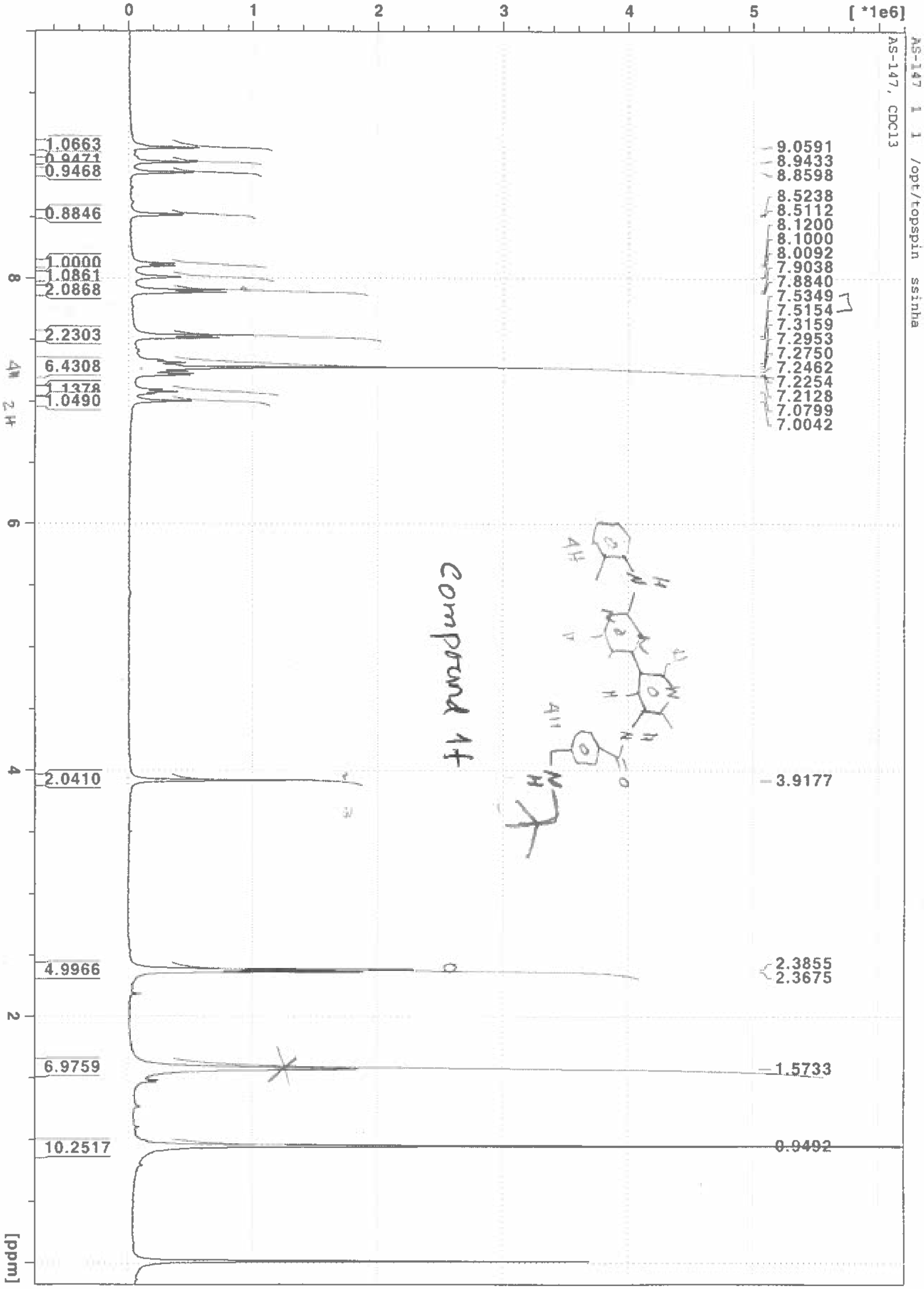
AS-147 1 1 /opt/topspin ssinha
AS-147, CDCl3
[ *1e6]
0
1
2
3
4
5
[ppm]
8
6
4
2
1.0663
0.9471
0.9468
0.8846
1.0000
1.0861
2.0868
2.2303
6.4308
1.1378
1.0490
2.0410
4.9966
6.9759
10.2517
9.0591
8.9433
8.8598
8.5238
8.5112
8.1200
8.1000
8.0092
7.9038
7.8840
7.5349
7.5154
7.3159
7.2953
7.2750
7.2462
7.2254
7.2128
7.0799
7.0042
3.9177
2.3855
2.3675
1.5733
0.9492

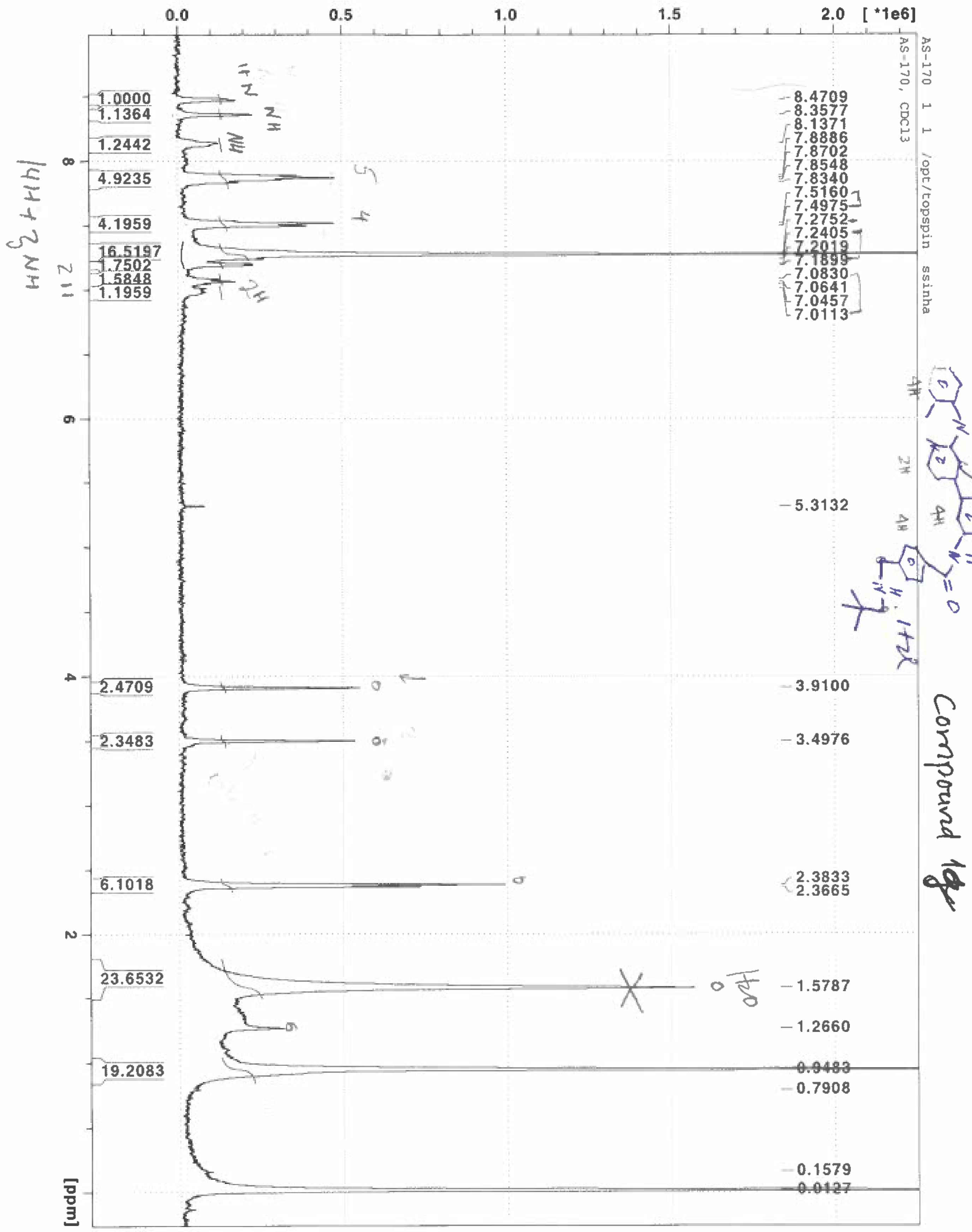
AS-170 1 1 /opt/topspin ssinha
AS-170, CDCl3
[*1e6]
0.0
0.5
1.0
1.5
2.0
8.4709
8.3577
8.1371
7.8886
7.8702
7.8548
7.8340
7.5160
7.4975
7.2752
7.2405
7.2019
7.1899
7.0830
7.0641
7.0457
7.0113
5.3132
3.9100
3.4976
2.3833
2.3665
1.5787
1.2660
0.8483
0.7908
0.1579
0.0127
1.0000
1.1364
1.2442
4.9235
4.1959
16.5197
1.7502
1.5848
1.1959
2.4709
2.3483
6.1018
23.6532
19.2083
8
6
4
2
[ppm]
Compound 1g

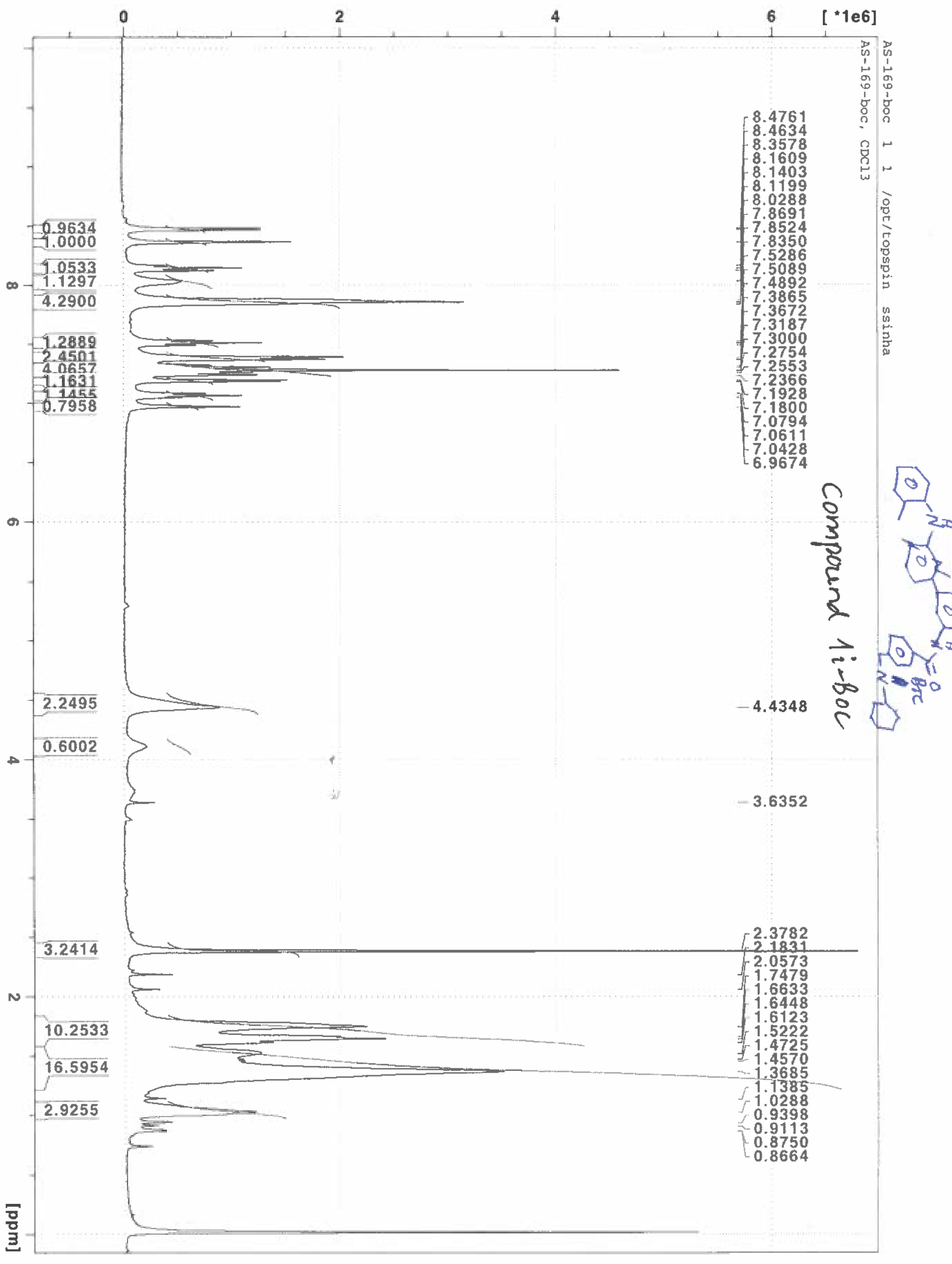
AS-169-boc 1 1 /opt/topspin ssinha
AS-169-boc, CDCl3
[ *1e6]
0
2
4
6
[ppm]
8
6
4
2
8.4761
8.4634
8.3578
8.1609
8.1403
8.1199
8.0288
7.8691
7.8524
7.8350
7.5286
7.5089
7.4892
7.3865
7.3672
7.3187
7.3000
7.2754
7.2553
7.2366
7.1928
7.1800
7.0794
7.0611
7.0428
6.9674
4.4348
3.6352
2.3782
2.1831
2.0573
1.7479
1.6633
1.6448
1.6123
1.5222
1.4725
1.4570
1.3685
1.1385
1.0288
0.9398
0.9113
0.8750
0.8664
0.9634
1.0000
1.0533
1.1297
4.2900
1.2889
2.4501
4.0657
1.1631
1.1455
0.7958
2.2495
0.6002
3.2414
10.2533
16.5954
2.9255
Compound 1i-Boc



1i-boc

AS-2-91 CDCl3

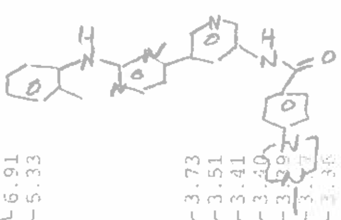

Compound 1j

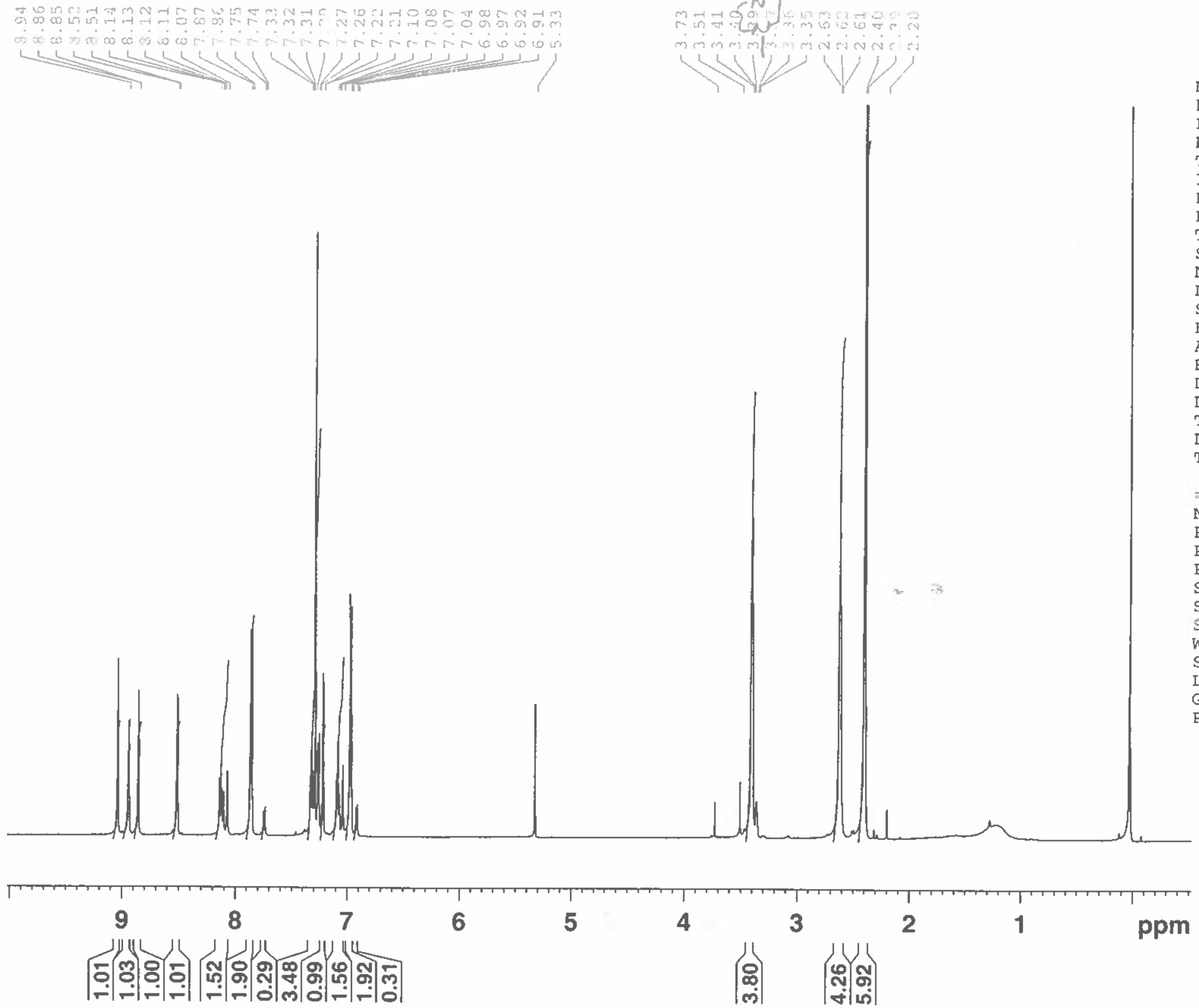


| | |
|---|---|
| NAME | AS-2-91 |
| EXPNO | 1 |
| PROCNO | 1 |
| Date_ | 20160211 |
| Time | 16.44 |
| INSTRUM | spect |
| PROBHD | 5 mm CPTCI 1H- |
| PULPROG | zg30 |
| TD | 32768 |
| SOLVENT | CDCl3 |
| NS | 16 |
| DS | 8 |
| SWH | 7788.162 Hz |
| FIDRES | 0.237676 Hz |
| AQ | 2.1038198 sec |
| RG | 28.5 |
| DW | 64.200 usec |
| DE | 6.00 usec |
| TE | 298.1 K |
| D1 | 1.00000000 sec |
| TD0 | 1 |

======== CHANNEL f1 ========

| | |
|---|---|
| NUC1 | 1H |
| P1 | 7.45 usec |
| PL1 | 4.50 dB |
| PL1W | 5.70400620 W |
| SFO1 | 600.1728538 MHz |
| SI | 16384 |
| SF | 600.1699972 MHz |
| WDW | EM |
| SSB | 0 |
| LB | 1.00 Hz |
| GB | 0 |
| PC | 1.00 |

AS-2-88-1 CDCl3

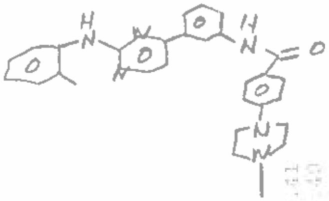

Compound 1k

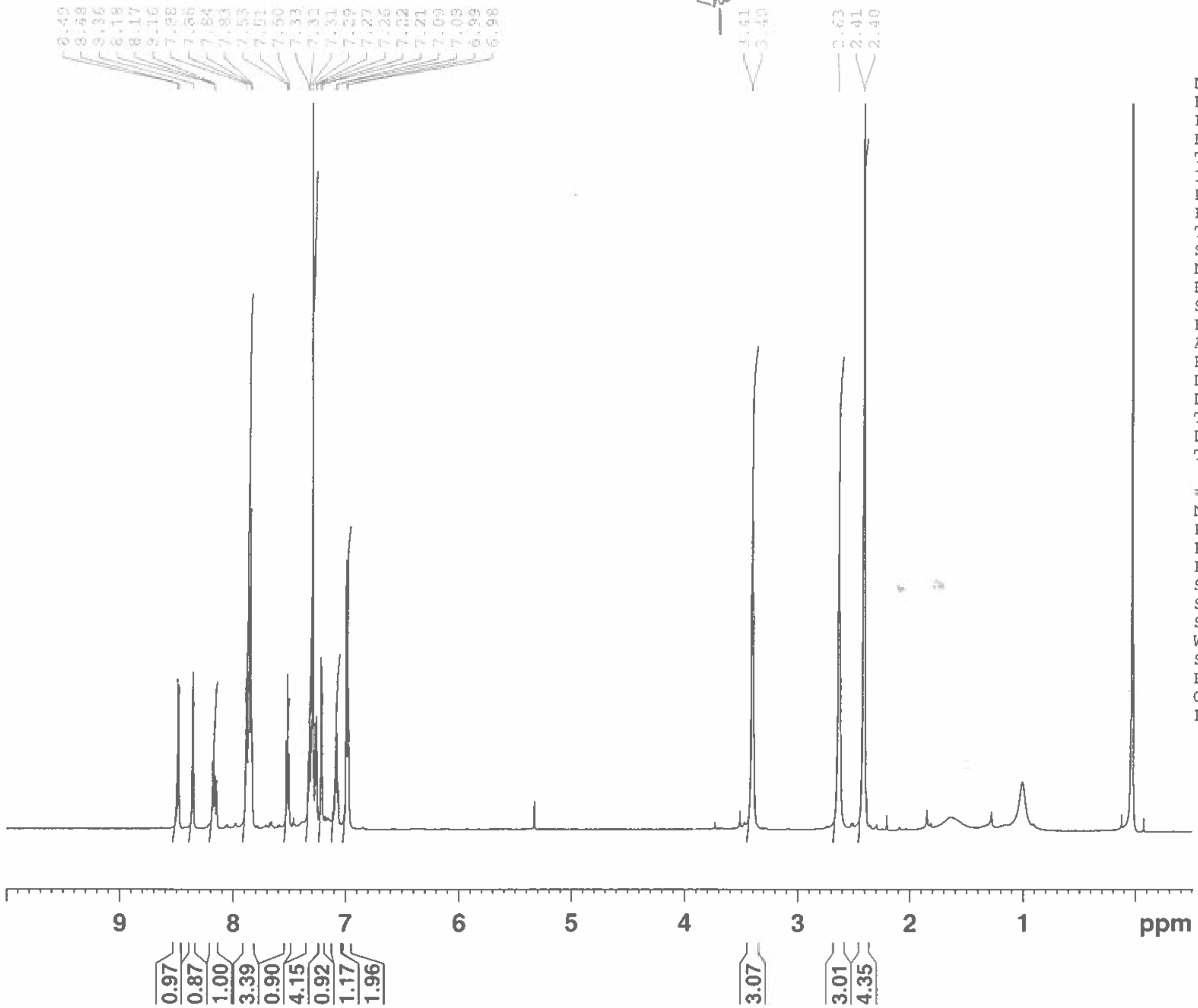


| | |
|---|---|
| NAME | AS-2-88-1 |
| EXPNO | 1 |
| PROCNO | 1 |
| Date_ | 20160211 |
| Time | 16.31 |
| INSTRUM | spect |
| PROBHD | 5 mm CPTCI 1H- |
| PULPROG | zg30 |
| TD | 32768 |
| SOLVENT | CDCl3 |
| NS | 16 |
| DS | 8 |
| SWH | 7788.162 Hz |
| FIDRES | 0.237676 Hz |
| AQ | 2.1038198 sec |
| RG | 28.5 |
| DW | 64.200 usec |
| DE | 6.00 usec |
| TE | 298.2 K |
| D1 | 1.00000000 sec |
| TD0 | 1 |

======== CHANNEL f1 ========

| | |
|---|---|
| NUC1 | 1H |
| P1 | 7.45 usec |
| PL1 | 4.50 dB |
| PL1W | 5.70400620 W |
| SFO1 | 600.1728538 MHz |
| SI | 16384 |
| SF | 600.1699972 MHz |
| WDW | EM |
| SSB | 0 |
| LB | 1.00 Hz |
| GB | 0 |
| PC | 1.00 |

AS-2-65

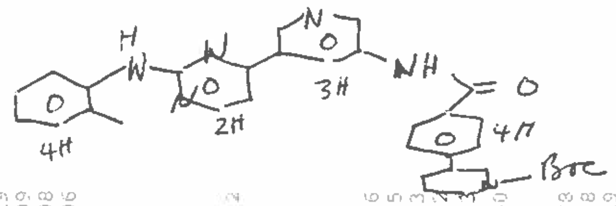


Compound 1l-Boc

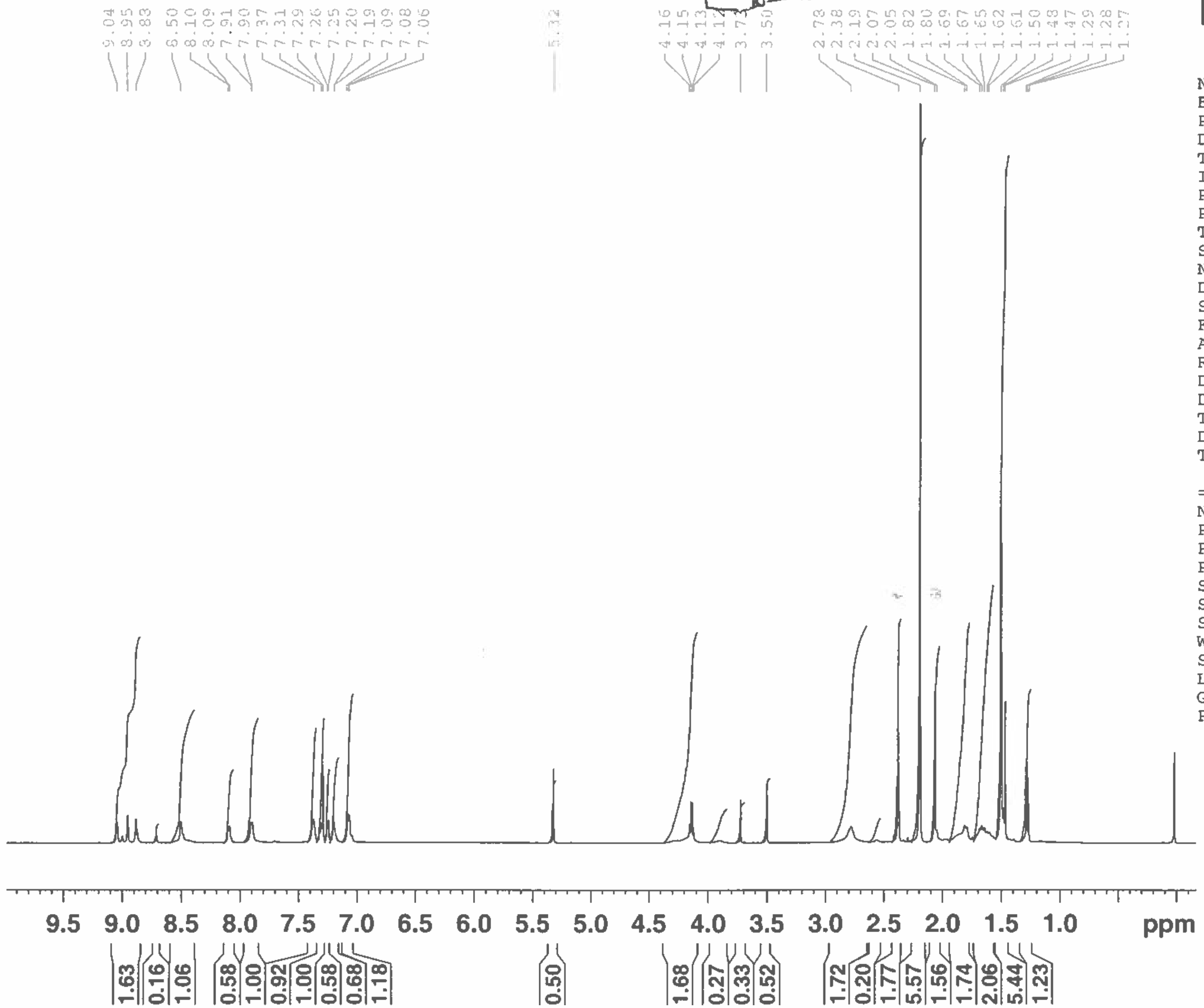


BRUKER

| | |
|---|---|
| NAME | AS-2-65 |
| EXPNO | 1 |
| PROCNO | 1 |
| Date_ | 20151008 |
| Time | 18.10 |
| INSTRUM | spect |
| PROBHD | 5 mm CPTCI 1H- |
| PULPROG | zg30 |
| TD | 32768 |
| SOLVENT | CDCl3 |
| NS | 16 |
| DS | 8 |
| SWH | 7788.162 Hz |
| FIDRES | 0.237676 Hz |
| AQ | 2.1038198 sec |
| RG | 12.7 |
| DW | 64.200 usec |
| DE | 6.00 usec |
| TE | 298.2 K |
| D1 | 1.00000000 sec |
| TD0 | 1 |

======== CHANNEL f1 ========

| | |
|---|---|
| NUC1 | 1H |
| P1 | 7.45 usec |
| PL1 | 4.50 dB |
| PL1W | 5.70400620 W |
| SFO1 | 600.1728538 MHz |
| SI | 16384 |
| SF | 600.1699972 MHz |
| WDW | EM |
| SSB | 0 |
| LB | 1.00 Hz |
| GB | 0 |
| PC | 1.00 |

New lot 1/8/16
AS-2-262-boc

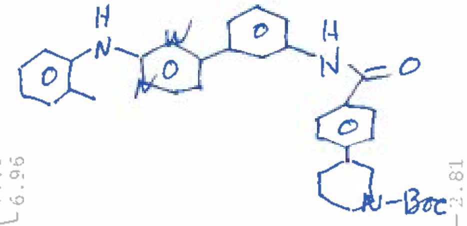

Compound 1m-Boc

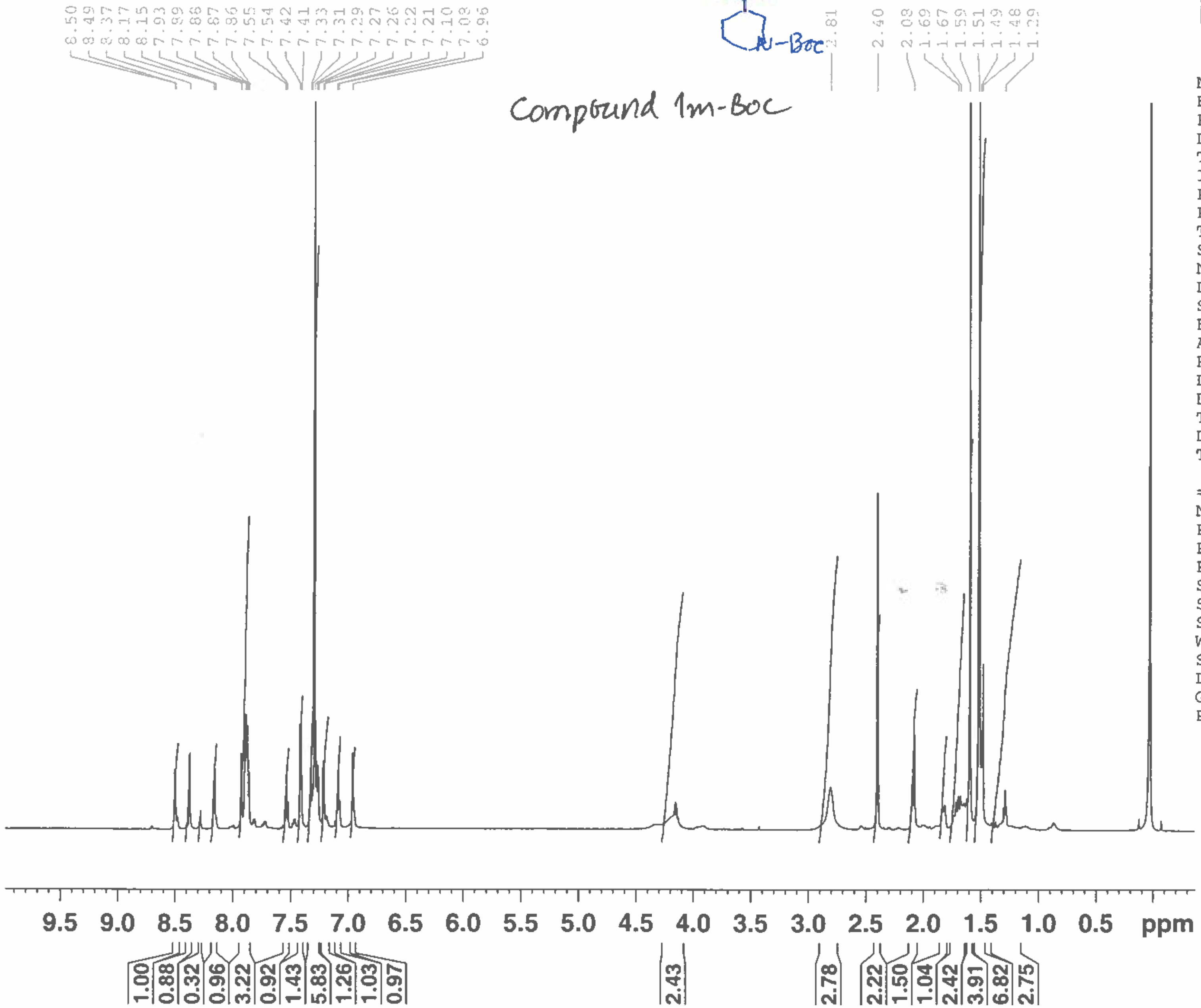


BRUKER

| | |
|---|---|
| NAME | AS-262-boc |
| EXPNO | 1 |
| PROCNO | 1 |
| Date_ | 20160108 |
| Time | 12.06 |
| INSTRUM | spect |
| PROBHD | 5 mm CPTCI 1H- |
| PULPROG | zg30 |
| TD | 32768 |
| SOLVENT | CDCl3 |
| NS | 16 |
| DS | 8 |
| SWH | 7788.162 Hz |
| FIDRES | 0.237676 Hz |
| AQ | 2.1038198 sec |
| RG | 20.2 |
| DW | 64.200 usec |
| DE | 6.00 usec |
| TE | 298.2 K |
| D1 | 1.00000000 sec |
| TD0 | 1 |

======== CHANNEL f1 ========

| | |
|---|---|
| NUC1 | 1H |
| P1 | 7.45 usec |
| PL1 | 4.50 dB |
| PL1W | 5.70400620 W |
| SFO1 | 600.1728538 MHz |
| SI | 16384 |
| SF | 600.1699972 MHz |
| WDW | EM |
| SSB | 0 |
| LB | 1.00 Hz |
| GB | 0 |
| PC | 1.00 |

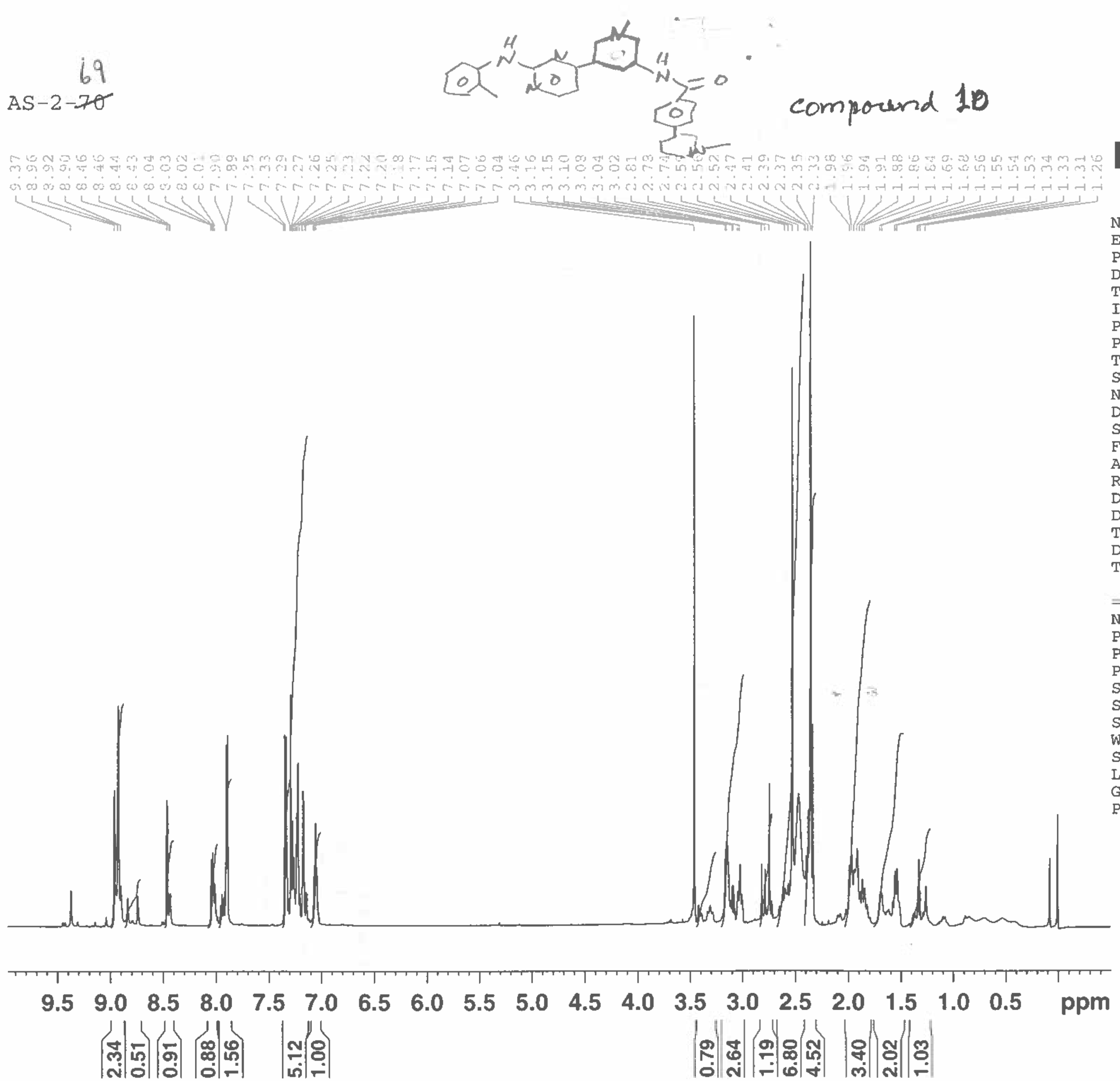


BRUKER

| | |
|---|---|
| NAME | AS-2-70 |
| EXPNO | 1 |
| PROCNO | 1 |
| Date_ | 20151026 |
| Time | 16.37 |
| INSTRUM | spect |
| PROBHD | 5 mm CPTCI 1H- |
| PULPROG | zg30 |
| TD | 32768 |
| SOLVENT | CDCl3 |
| NS | 16 |
| DS | 8 |
| SWH | 7788.162 Hz |
| FIDRES | 0.237676 Hz |
| AQ | 2.1038198 sec |
| RG | 12.7 |
| DW | 64.200 usec |
| DE | 6.00 usec |
| TE | 298.2 K |
| D1 | 1.00000000 sec |
| TD0 | 1 |

======== CHANNEL f1 ========

| | |
|---|---|
| NUC1 | 1H |
| P1 | 7.45 usec |
| PL1 | 4.50 dB |
| PL1W | 5.70400620 W |
| SFO1 | 600.1728538 MHz |
| SI | 16384 |
| SF | 600.1699972 MHz |
| WDW | EM |
| SSB | 0 |
| LB | 1.00 Hz |
| GB | 0 |
| PC | 1.00 |

AS-2-69

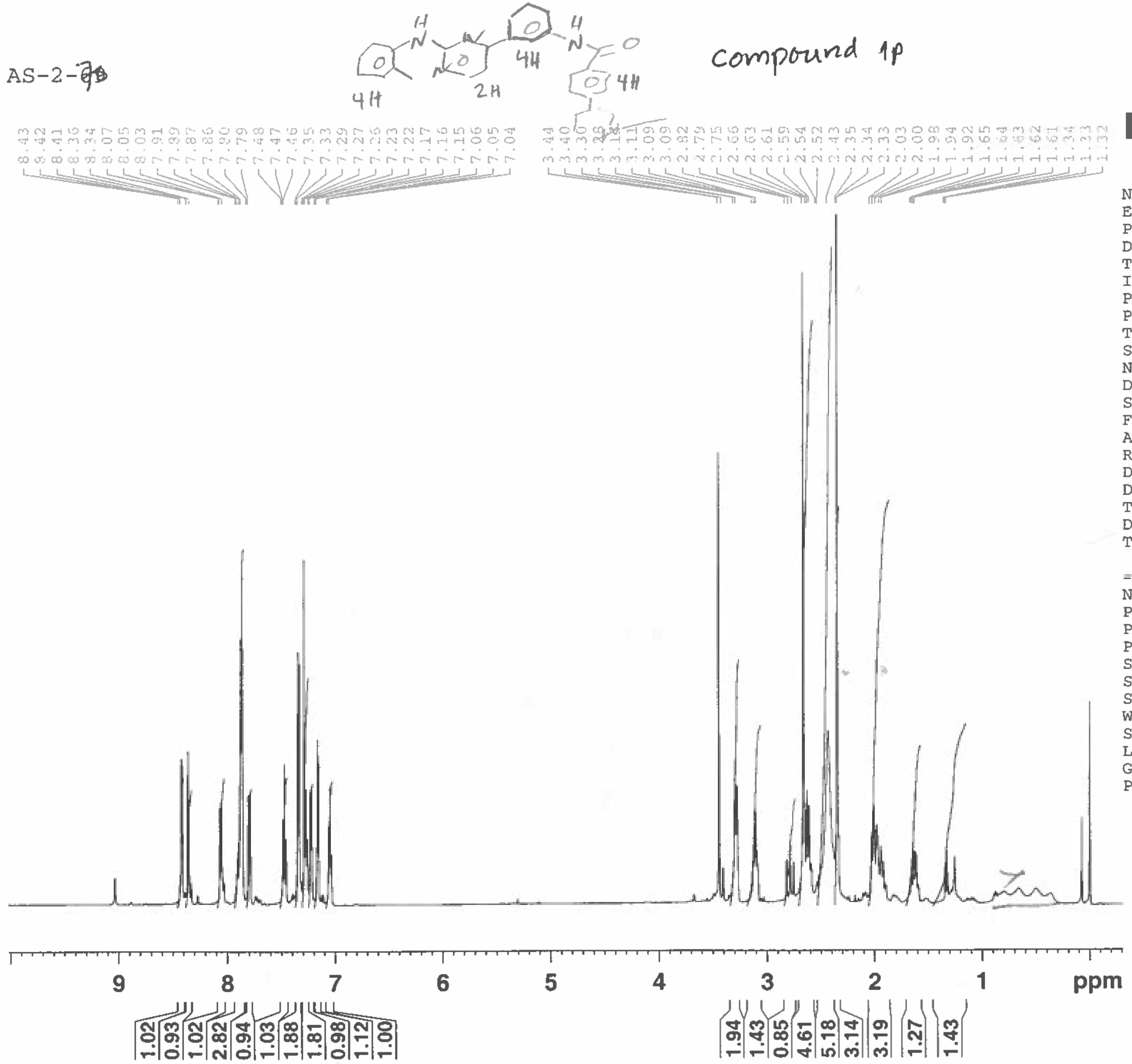


| | |
|---|---|
| NAME | AS-2-69 |
| EXPNO | 1 |
| PROCNO | 1 |
| Date_ | 20151026 |
| Time | 16.42 |
| INSTRUM | spect |
| PROBHD | 5 mm CPTCI 1H- |
| PULPROG | zg30 |
| TD | 32768 |
| SOLVENT | CDCl3 |
| NS | 16 |
| DS | 8 |
| SWH | 7788.162 Hz |
| FIDRES | 0.237676 Hz |
| AQ | 2.1038198 sec |
| RG | 12.7 |
| DW | 64.200 usec |
| DE | 6.00 usec |
| TE | 298.2 K |
| D1 | 1.00000000 sec |
| TD0 | 1 |

======== CHANNEL f1 ========

| | |
|---|---|
| NUC1 | 1H |
| P1 | 7.45 usec |
| PL1 | 4.50 dB |
| PL1W | 5.70400620 W |
| SFO1 | 600.1728538 MHz |
| SI | 16384 |
| SF | 600.1699972 MHz |
| WDW | EM |
| SSB | 0 |
| LB | 1.00 Hz |
| GB | 0 |
| PC | 1.00 |

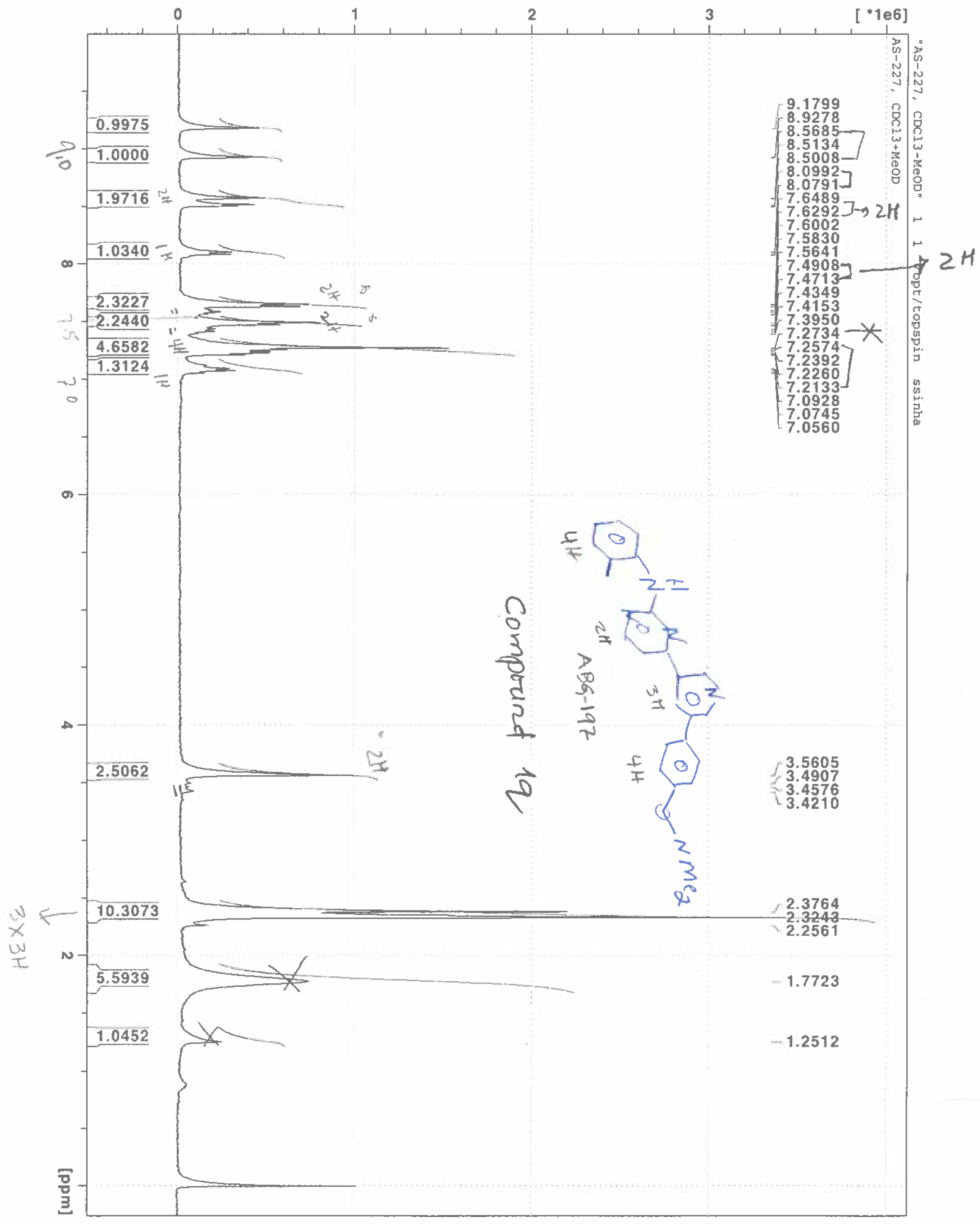

"AS-227, CDCl3-MeOD" 1 1 "/opt/topspin ssinha"
AS-227, CDCl3+MeOD
[ *1e6]
0
1
2
3
9.1799
8.9278
8.5685
8.5134
8.5008
8.0992
8.0791
7.6489
7.6292
7.6002
7.5830
7.5641
7.4908
7.4713
7.4349
7.4153
7.3950
7.2734
7.2574
7.2392
7.2260
7.2133
7.0928
7.0745
7.0560
3.5605
3.4907
3.4576
3.4210
2.3764
2.3243
2.2561
1.7723
1.2512
0.9975
1.0000
1.9716
1.0340
2.3227
2.2440
4.6582
1.3124
2.5062
10.3073
5.5939
1.0452
8
6
4
2
0
[ppm]

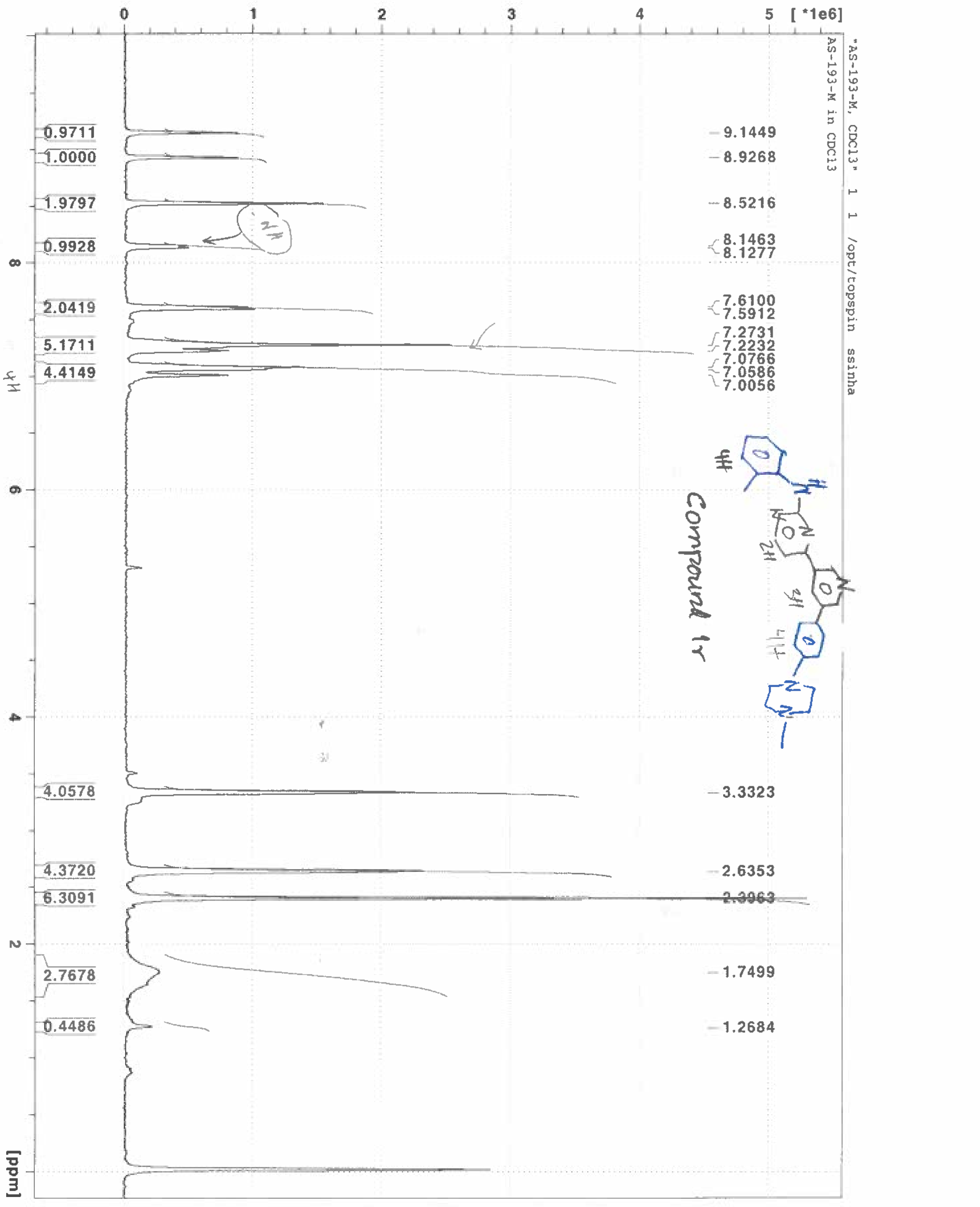
"AS-193-M, CDCl3" 1 1 /opt/topspin ssinha
AS-193-M in CDCl3
[ *1e6]
0 1 2 3 4 5
9.1449
8.9268
8.5216
8.1463
8.1277
7.6100
7.5912
7.2731
7.2232
7.0766
7.0586
7.0056
3.3323
2.6353
2.3963
1.7499
1.2684
0.9711
1.0000
1.9797
0.9928
2.0419
5.1711
4.4149
4.0578
4.3720
6.3091
2.7678
0.4486
8
6
4
2
[ppm]
Compound 1r

AS-2-117

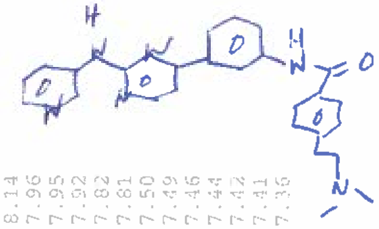

Compound 1s

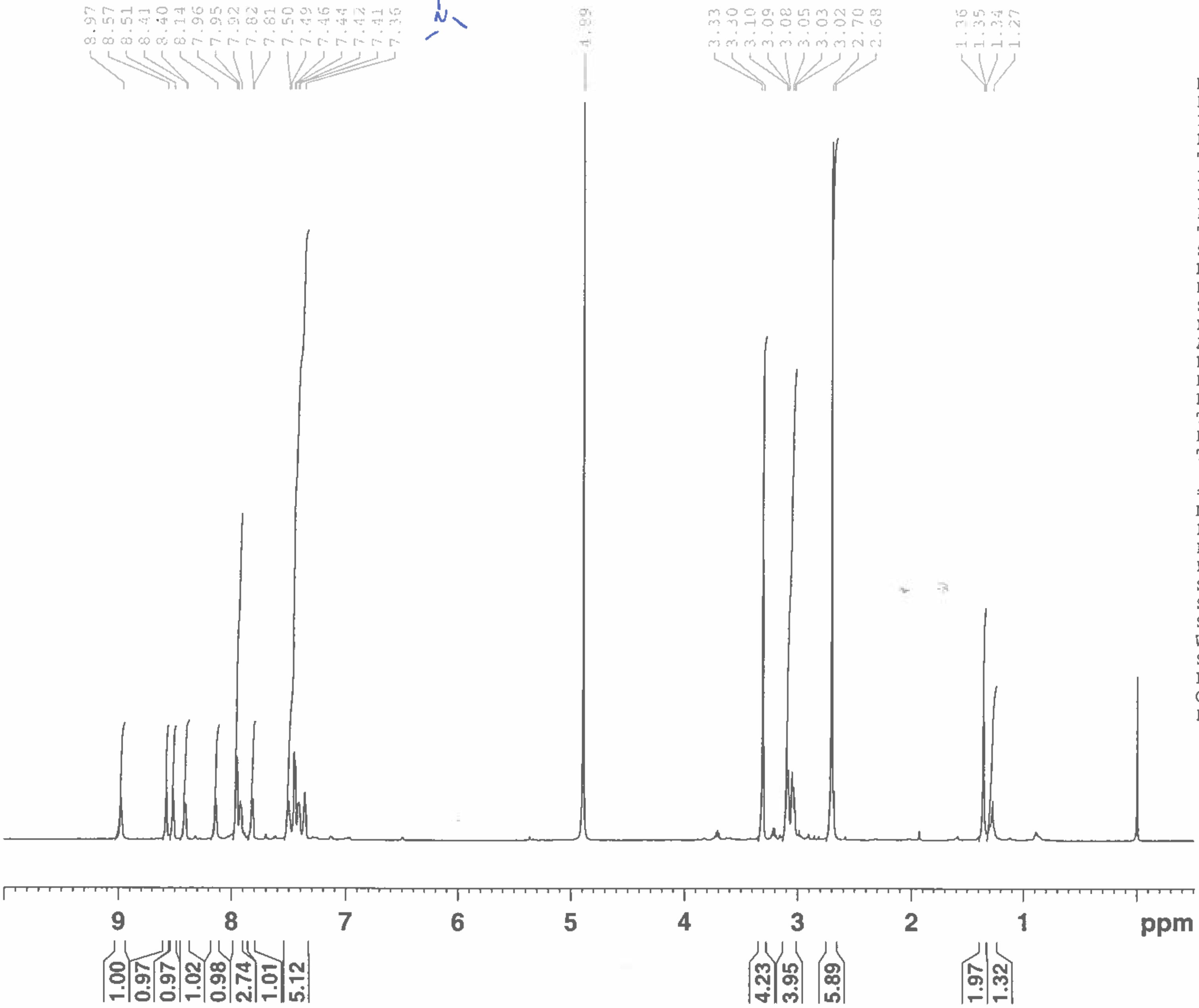


BRUKER

| | |
|---|---|
| NAME | AS-2-117 |
| EXPNO | 1 |
| PROCNO | 1 |
| Date_ | 20160428 |
| Time | 16.09 |
| INSTRUM | spect |
| PROBHD | 5 mm CPTCI 1H- |
| PULPROG | zg30 |
| TD | 32768 |
| SOLVENT | MeOD |
| NS | 16 |
| DS | 8 |
| SWH | 7788.162 Hz |
| FIDRES | 0.237676 Hz |
| AQ | 2.1038198 sec |
| RG | 16 |
| DW | 64.200 usec |
| DE | 6.00 usec |
| TE | 298.1 K |
| D1 | 1.00000000 sec |
| TD0 | 1 |

======== CHANNEL f1 ========

| | |
|---|---|
| NUC1 | 1H |
| P1 | 7.45 usec |
| PL1 | 4.50 dB |
| PL1W | 5.70400620 W |
| SFO1 | 600.1728538 MHz |
| SI | 16384 |
| SF | 600.1700169 MHz |
| WDW | EM |
| SSB | 0 |
| LB | 1.00 Hz |
| GB | 0 |
| PC | 1.00 |

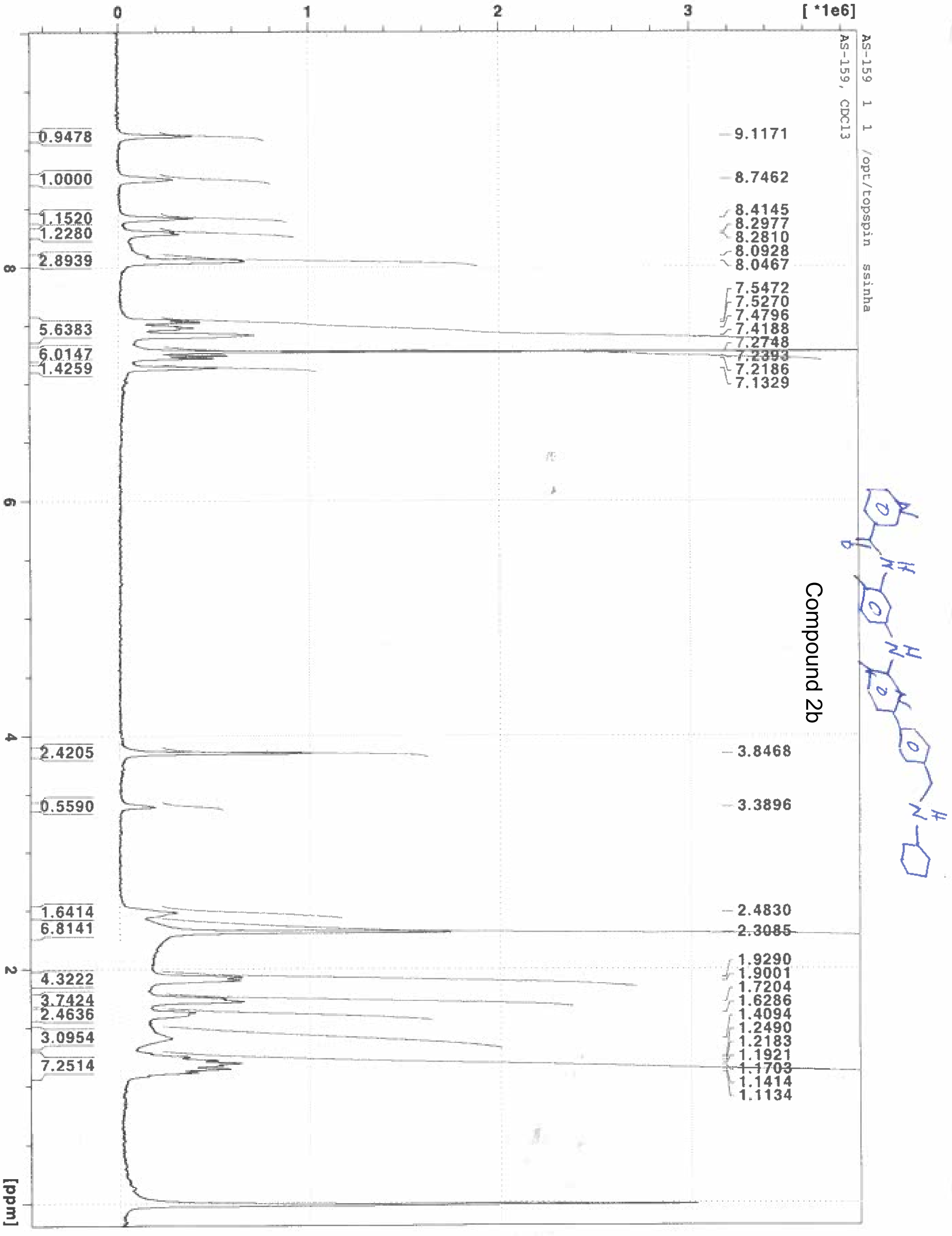
AS-159 1 1 /opt/topspin ssinha
AS-159, CDCl3
[ *1e6]
[ppm]
9.1171
8.7462
8.4145
8.2977
8.2810
8.0928
8.0467
7.5472
7.5270
7.4796
7.4188
7.2748
7.2393
7.2186
7.1329
3.8468
3.3896
2.4830
2.3085
1.9290
1.9001
1.7204
1.6286
1.4094
1.2490
1.2183
1.1921
1.1703
1.1414
1.1134
0.9478
1.0000
1.1520
1.2280
2.8939
5.6383
6.0147
1.4259
2.4205
0.5590
1.6414
6.8141
4.3222
3.7424
2.4636
3.0954
7.2514

Compound 2b

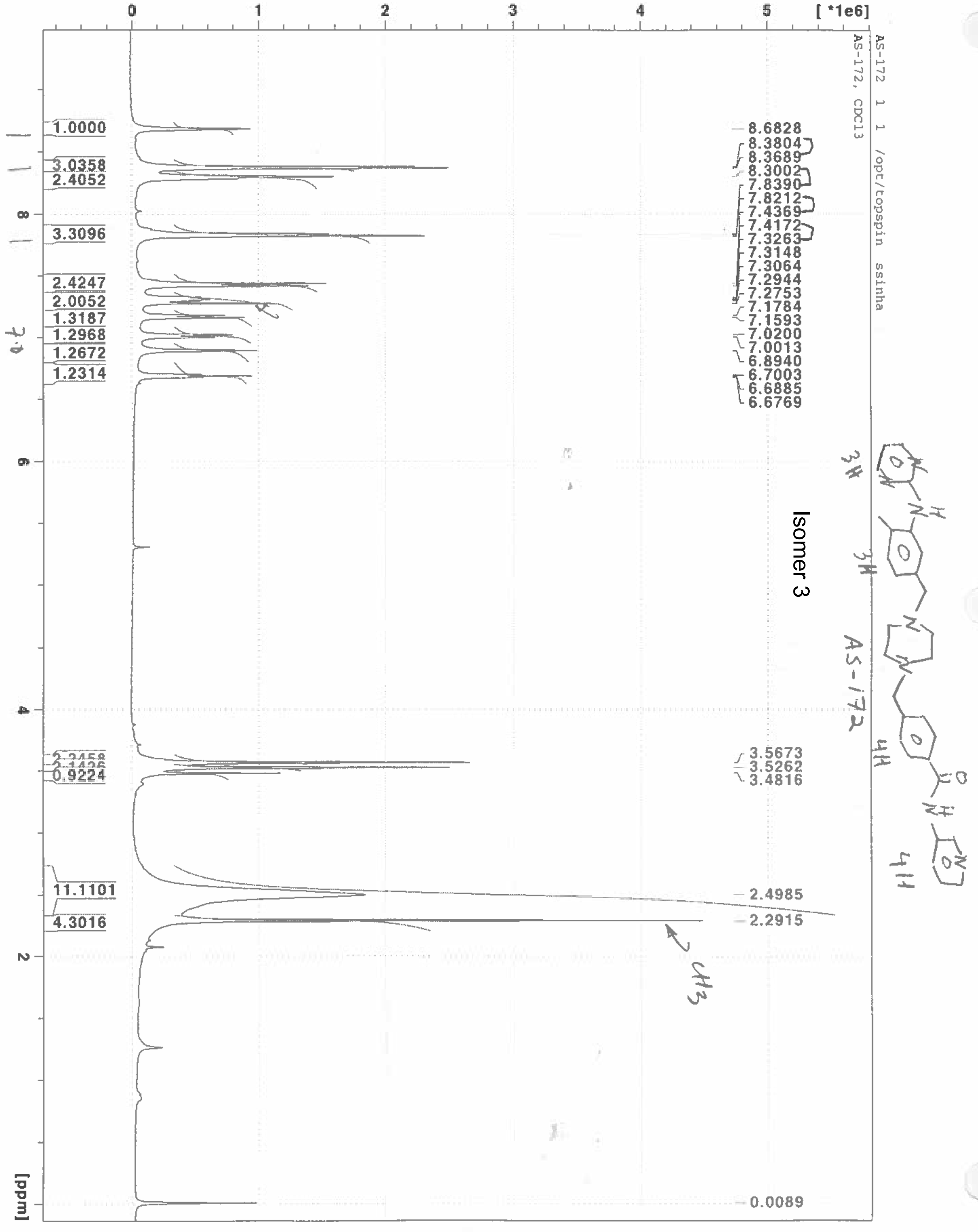
AS-172 1 1 /opt/topspin ssinha
AS-172, CDCl3
Isomer 3
[ *1e6]
[ppm]
8.6828
8.3804
8.3689
8.3002
7.8390
7.8212
7.4369
7.4172
7.3263
7.3148
7.3064
7.2944
7.2753
7.1784
7.1593
7.0200
7.0013
6.8940
6.7003
6.6885
6.6769
3.5673
3.5262
3.4816
2.4985
2.2915
0.0089
1.0000
3.0358
2.4052
3.3096
2.4247
2.0052
1.3187
1.2968
1.2672
1.2314
0.9224
11.1101
4.3016
